# Sewing the future of cotton: a multi-omics study combining nanomechanics, transcriptomics, and phenotypic traits

**DOI:** 10.1101/2025.05.12.653486

**Authors:** Md Hasibul Hasan Hasib, Nabila Masud, Angona Biswas, Talukder Zaki Jubery, Carson Stanley, Sivakumar Swaminathan, Corrinne E. Grover, Jonathan F. Wendel, Soumik Sarkar, Olga A. Zabotina, Anwesha Sarkar

## Abstract

Cellulose microfibrils that are essential for mechanical strength and overall quality of cotton fibers. This study quantifies and compares the nanoscale structural and mechanical properties of cellulose microfibrils such as microfibril dimensions, crossover count and angles, roughness, and Young’s modulus for two popular cotton species: *Gossypium hirsutum* (Gh) and *Gossypium barbadense* (Gb) fibers across four growth stages (8, 12, 18, and 22 days post-anthesis) using atomic force microscopy (AFM). Our results revealed for the first time that Gb fibers exhibit a better alignment, finer dimensions, and higher stiffness compared to Gh fibers at nanoscale, resulting in smoother fiber surfaces, and improved quality at macroscale. We are also the first to develop machine-learning models to predict macroscale phenotypic traits specifically boll length and cellulose content using nanoscale features alone and in combination with multi-omics modalities, substantially enhancing the predictive accuracy and highlighting opportunities for robust cross-species modeling of cotton fiber traits.

---

Cotton fibers from *Gossypium* species are the most important natural fibers for the textile industry worldwide. Among the cultivated species, *Gossypium hirsutum* (Gh) inherits mid-quality fiber, but it is extensively cultivated worldwide due to its high productivity and environmental adaptability. In contrast, *Gossypium barbadense* (Gb) inherits better fiber quality, characterized by greater length, fineness, and strength [1, 2]. Cotton fibers are individual single cells that emerge from the seed coat epidermis and can grow more than 2.5-4 cm in length. The mature cotton fiber is made up of developed cell walls. These cell wall constituents are polysaccharides, such as pectin, hemicellulose, and cellulose. Their dynamic rearrangements during fiber growth regulate fiber quality [3]. Investigating the nanoscale structural, topographical and mechanical properties of the fiber cell wall throughout the developmental stages could provide valuable insights into the macroscale differences in the cotton fiber quality. Mature fibers contain more than 95% cellulose, which is made up of chains of glucose molecules. These chains aggregate and form microfibrils [4]. Cellulose is biosynthesized by the plasma membrane-localized cellulose synthase (CESA) complex. CESA1, CESA3, and CESA6 synthesize primary cell wall (PCW) cellulose, while CESA4, CESA7, and CESA8 are involved in synthesizing secondary cell wall (SCW) cellulose [5]. Upon fertilization (anthesis), the cotton fiber cells bud out from the seed coat epidermis. Their development occurs through the fiber cells growth, and development consists of five overlapping stages, namely, initiation, elongation, transition, SCW deposition, and maturation [6, 7]. During the initiation phase (0–2 DPA), epidermal cells differentiate into fiber initials. This is followed by the elongation phase (3–16 DPA), during which fibers undergo rapid anisotropic growth, facilitated by the dynamic remodeling of the PCW and the transverse alignment of cellulose microfibrils, allowing directional expansion [8–10]. The transition phase (16–20 DPA) is marked by the cessation of PCW fiber elongation and the beginning of SCW cellulose accumulation. The SCW deposition phase (20–35 DPA) involves heavy accumulation of cellulose, and organization of cellulose into densely packed, helically arranged microfibrils, which enhance the mechanical robustness, strength, and stiffness of fiber, and eventually, leads to drying, twisting, and maturation of fiber (35-50 DPA) [3, 11, 12]. These developmental phases collectively determine the fiber’s length, strength, and quality, with the hierarchical organization of microfibrils being a critical factor. Previous molecular studies [13, 14] have revealed significant differences in the expression of genes involved in the biosynthesis and dynamic remodeling of cellulose, pectin, and xyloglucan polysaccharides in the cell wall during the critical growth stages between Gh and Gb. Keynia and Sedighe et al. [15] investigated the role of cellulose microfibril arrangement in tensile strength and mechanical performance. Machine learning (ML) [16] has also been used to predict complex traits from gene expression data in crops like cotton. Liu et al. [17] showed that models based on transcript levels of fiber length genes can accurately predict cotton fiber length. Similarly, Khalilisamani et al. [18] demonstrated that integrating gene expression with single-nucleotide polymorphisms (SNP) data through gene network analysis improved prediction accuracy for fiber traits such as elongation and strength. In other crops:

*Arabidopsis*, maize, and rice – ML models using multiomics data have also improved trait prediction. Wang et al. [19] combined genomic, transcriptomic, and methylomic data to predict traits in *Arabidopsis thaliana*. These studies support using ML-based multiomics approaches for precise phenotype prediction and efficient genotype selection in crop breeding.

However, the nanoscale structural, topographical, and mechanical properties of these microfibrils: *Nano-omics* remain largely unexplored in these studies. The variations in these nanoscale properties across different cotton species are not well understood. Therefore, we address this current knowledge gap by analyzing structural and nanomechanical properties of cell walls at the single-molecule level using bio-atomic force microscopy (Bio-AFM). Unlike Scanning Electron Microscopy (SEM) and Transmission Electron Microscopy (TEM), which require extensive sample preparation that can distort fibers, AFM [20–31] preserves their structural integrity while mapping both topographical and mechanical properties. In our previous study [25], we used AFM to investigate and analyze the nanoscale properties of cellulose microfibrils of *Arabidopsis thaliana* in wild-type (WT) and mutant populations.

This study aims to conduct a novel, comprehensive multiscale characterization of Gh and Gb cotton fibers by integrating nanomechanical and macrostructural analyses. Using AFM in PeakForce QNM air mode, we quantified key nanoscale parameters such as microfibril orientation, crossover count, angles, roughness, Young’s modulus, diameter, and height during critical growth stages 8, 12, 18, and 22 days postanthesis (DPA). These nanoscale findings were correlated with macro-level traits such as cellulose deposition and cotton boll size to determine factors influencing fiber quality. We developed a machine learning approach that leverages multi-omics data by integrating nanomechanical properties (obtained from AFM) with transcriptomic data (from CESA genes) to predict fiber phenotypes in both Gh and Gb. We compared models built on combined AFM and CESA features to those using only AFM or CESA features. Additionally, we quantified the contribution of each feature and evaluated model performance by training on Gh data to predict Gb fiber traits. These findings provide a framework for breeding programs, supporting the development of biotechnological strategies to enhance cotton fiber quality without compromising yield.

## Results

### Role of Cellulose Content in Cotton Fiber Development

Cellulose is primarily responsible for strength, flexibility, and durability of cotton fibers. It forms the backbone of the SCW, determining the fiber’s ability to withstand mechanical stress and its quality in textile production. Fig. 1(a) presents the cellulose content for both cotton species demonstrating minimal accumulation at the beginning (up to 16 DPA for Gh and up to 19 DPA for Gb) and this phase emphasizes fiber elongation by growth rate and PCW formation. After 16 DPA in Gh and 19 DPA in Gb, cellulose deposition rapidly increases, coinciding with the SCW synthesis stage. Gh fibers display faster cellulose accumulation, reflecting an earlier transition to SCW thickening, leading to bulkier cotton bolls (Supplementary Fig. S1(d)). Conversely, Gb shows slower cellulose deposition but lighter cotton bolls than Gh. There is also a distinction in boll size between Gh and Gb, as shown in Fig. 1(b, c). By comparing the four growth stages (8, 12, 18, and 22 DPA) of both species, Gb tends to produce longer bolls due to its extended growth phase and prolonged elongation.

### Dynamic Microfibril Architecture: Comparative Analysis

The crossover count (the number of intersections between adjacent microfibrils) and the angles between these intersections are key indicators of fiber organization. Fig. 2(a-h) presents high-resolution Bio-AFM images, showcasing the crossover count, angles between adjacent microfibrils, and intensity fluctuations along individual microfibrils. These images show that at earlier stages (8 and 12 DPA), crossover frequency and angles are high, whereas at later stages (18 and 22 DPA), they decrease significantly, with microfibrils becoming nearly parallel. At 8 and 12 DPA (Fig. 2(i,j)), both cotton species exhibit a high crossover count per 500 × 500 nm^2^ areas (Gh 8DPA: 37.5; Gh 12 DPA: 17.8 and Gb 8 DPA: 41.3; Gb 12 DPA: 27.5) with interception angles (Gh 8DPA: 73.3°; Gh 12 DPA: 60.5° and Gb 8 DPA: 71.2°; Gb 12 DPA: 55.6°) that yield a less ordered PCW structure during the elongation phase. However, as development progresses, both species show a significant reduction in crossover count at 18 and 22 DPA (Supplementary Fig. S1(a,b)), leading to more aligned microfibrils exhibiting narrower angles. This alignment can be attributed to the transition to SCW synthesis, where ordered cellulose deposition dominates. The increased microfibril density during cell wall synthesis indicates a more compact and cohesive architecture in Gb fibers compared to Gh. The intensity variations in the peak force error images may reflect the differences in microfibrils architectures and insight on cellulose deposition. At 8 and 12 DPA, both species display negligible cellulose content intensity patterns, corresponding to minimal cellulose distribution during the elongation phase. In contrast, 18 and 22 DPA fibers show more distinct intensity levels along their microfibrils (Gh 18DPA: 70 a.u.; Gh 22 DPA: 79.0 a.u. and Gb 18 DPA: 75.9 a.u.; Gb 22 DPA: 81.4 a.u.), indicating higher cellulose deposition and well-organized fiber structure (see Supplementary Fig. S1(c)).

### Surface Nanomechanics: Roughness and Young’s Modulus Comparison

The surface of microfibrils influences fiber quality, with smoother surfaces (fineness) and higher stiffness being indicators of superior fibers. At 8 and 12 DPA (Fig. 2(k)), the roughness values of both Gh and Gb are relatively low, reflecting the elongation phase, where cellulose deposition is minimal. The differences between the two species are minor at this stage, with Gb showing slightly smoother surfaces (0.33 nm and 0.52 nm for Gb vs. 0.50 nm and 0.55 nm for Gh at 8 and 12 DPA, respectively). As development progresses, the roughness increases significantly due to cellulose deposition during the SCW synthesis phase. At 18 and 22 DPA (Fig. 2(k)), Gh exhibits noticeably higher roughness compared to Gb (1.75 nm and 2.54 nm for Gh vs. 1.19 nm and 2.13 nm for Gb at 18 and 22 DPA, respectively). Also, Young’s modulus (a measure of fiber strength and stress tolerance) increases steadily for both species as the fibers mature and SCW deposit progresses (Fig. 2(l)). At 8 and 12 DPA, Gb demonstrates higher stiffness compared to Gh (68.9 MPa and 70.8 MPa for Gb vs. 55.1 MPa and 59.1 MPa for Gh at 8 and 12 DPA, respectively). This early advantage suggests a better initial fiber structure in Gb. By 18 and 22 DPA, the Young’s modulus for both species increases, reflecting the thickening and crystallization of the cellulose in the SCW. However, Gb consistently has a higher Young’s modulus than Gh (85.5 MPa and 87.2 MPa for Gb vs. 72.2 MPa and 83.5 MPa for Gh at 18 and 22 DPA, respectively), indicating a more mechanically robust fiber structure. Detailed comparisons among all

**Fig. 1.**
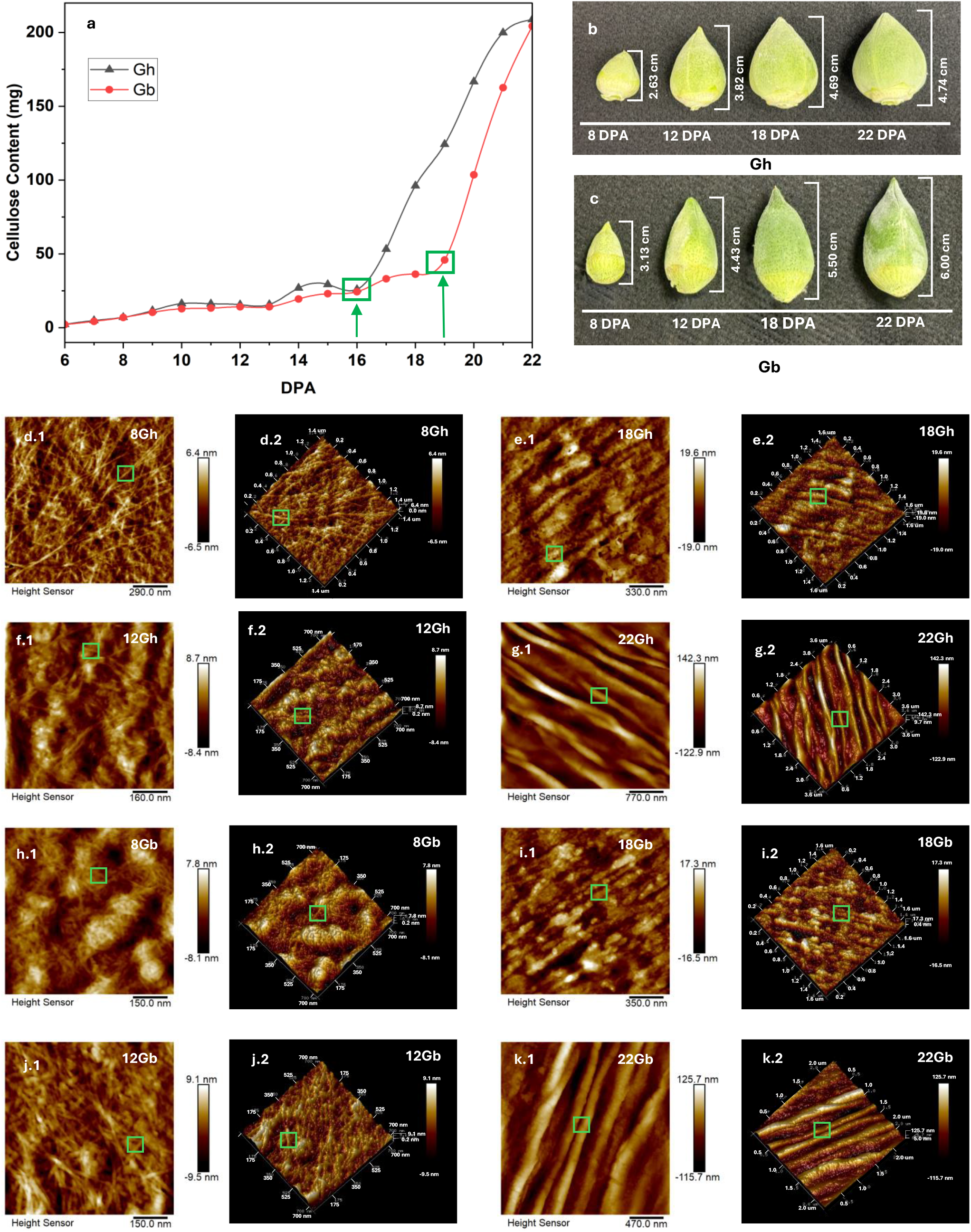
Comparative analysis of macroscale and nanoscale properties of *Gossypium hirsutum* (Gh) and *Gossypium barbadense* (Gb) across growth stages (8, 12, 18 and 22 DPA) (a) Cellulose content: reveals a sharp rise after 16 DPA in Gh and after 19 DPA in Gb that indicates differences in fiber elongation time. (b, c) Boll size progression of Gh and Gb from 8 to 22 DPA. (d1–k1) AFM microfibril height images and (d2–k2) corresponding 3D images provide nanoscale visualization of the microfibril structure. (d1, d2) Gh at 8 DPA, (f1, f2) Gh at 12 DPA, (e1, e2) Gh at 18 DPA, (g1, g2) Gh at 22 DPA, (h1, h2) Gb at 8 DPA, (j1, j2) Gb at 12 DPA, (i1, i2) Gb at 18 DPA, and (k1, k2) Gb at 22 DPA. Green boxes help to indicate single microfibrils as an example where the nanoscale measurements of individual microfibrils were analyzed.

**Fig. 2.**
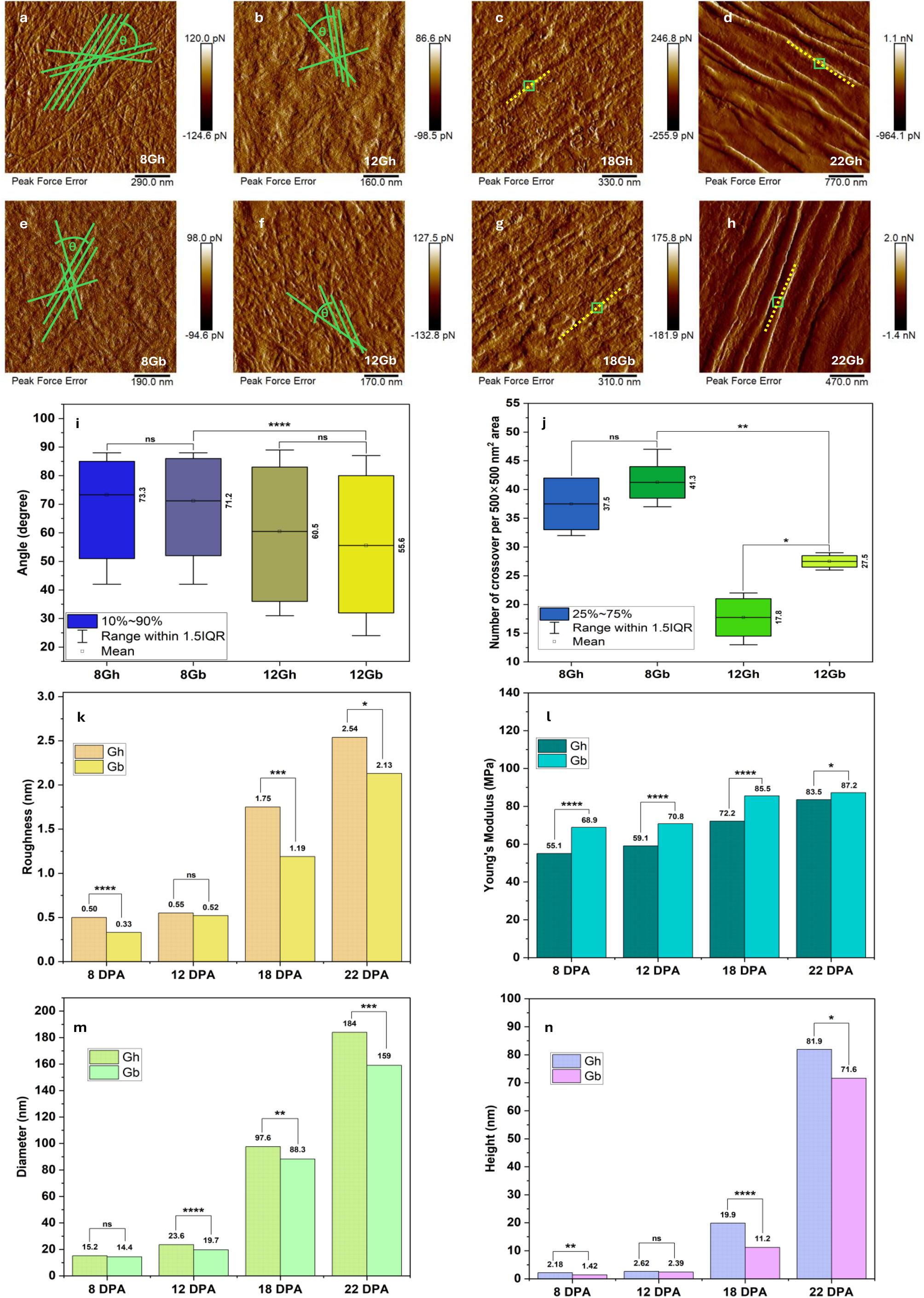
Comparative analysis of microfibrils properties in *Gossypium hirsutum* (Gh) and *Gossypium barbadense* (Gb) during cotton fiber development(8, 12, 18, and 22 DPA) (a–h) AFM peak force error images displaying microfibrils morphological structures in Gh (a–d) and Gb (e–h) at 8, 12, 18, and 22 DPA. Marked annotations visualize how crossover numbers, angles (a, b, e, f), and intensity variations (c, d, g, h) were quantitatively analyzed. (i, j) Quantification of microfibril crossover angles and crossover density (number of crossovers per 500 × 500 nm^2^ area), revealing structural differences between the two species at earlier growth stages 8 and 12 DPA. (k, l) Surface roughness (RMS) and Young’s modulus measurements highlight Gh fibers as consistently rougher, whereas Gb fibers exhibit smoother surfaces and higher mechanical strength across all growth stages (8, 12, 18, and 22 DPA). (m, n) Microfibril diameter and height measurements across different growth stages. Gh exhibits larger microfibril dimensions, whereas Gb shows smaller, more compact fibrils. Statistical significance is denoted as follows: ns indicates no significance, * represents p *<* 0.05, ** represents p *<* 0.01, *** represents p *<* 0.001, and **** corresponds to p *<* 0.0001. _5_ 8, 12, 18, and 22 DPA are in Supplementary Fig. S2(a, b, c, d).

### Nanoscale Microfibril Dimensions and Structural Comparisons

The diameter and height of microfibrils are key indicators of the structural development of cotton fibers. Fig. 2(m, n) quantifies microfibrils dimensions, showing a gradual increase in diameter and height for both Gh and Gb throughout the growth stages. At 8 and 12 DPA, Gh and Gb are narrower with smaller diameters (Gh:15.2 nm and 23.6 nm vs. Gb: 14.4 nm and 19.7 nm, respectively) and heights (Gh: 2.18 nm and 2.62 nm vs. Gb: 1.42 nm and 2.39 nm, respectively). By 18 and 22 DPA, Gh fibers are wider (97.6 nm and 184 nm vs. 88.3 nm and 159 nm, respectively), while heights are larger (Gh: 19.9 nm and 81.9 nm vs. Gb: 11.2 nm and 71.6 nm, respectively). These results indicate that Gh generally forms bulkier fibers and cotton bolls than Gb. Furthermore, Fig. 1(d-k) presents high-resolution Bio-AFM height images and corresponding 3D views of microfibrils at different growth stages for both cotton species. The green-boxed regions highlight individual microfibril, while height sensor and 3D images reveal their dimensional growth across DPA. Also, at 22 DPA, a twisting pattern emerges in single microfibril that might enhance the overall fiber strength. Detailed comparison among all 8, 12, 18, and 22 DPA for Gh and Gb are in Supplementary Fig. S3(a, b, c, d).

### Integrative Analysis Using Machine Learning

We evaluate the predictive performance of these nanoscale features in conferring macroscale phenotypic traits, i.e., boll length and cellulose content. Predictive performance is conducted using a 5-fold cross-validation with three different feature sets: (1) transcriptomics (CESAs-only); (2) nanomechanics (AFM-only); and (3) combined transcriptomics + nanomechanics (CESAs+AFM). The AFM dataset contains multiple microfibril measurements (width, height, roughness, Young’s modulus, angle, and crossover number), while the transcriptomics dataset (CESAs) contains gene expression measurements for the primary and secondary wall CESA-encoding genes (Supplementary Table 1-4). For both boll length: a proxy for fiber length (Fig. 3(a)) and cellulose content (Fig. 3(b)), the combined feature set: (CESAs+AFM) consistently achieves higher *R*^2^ values across all five folds compared with either CESAs-only or

**Table 1.** Predictive Performance of Regression Models Using Combined AFM and CESAs Features. This table presents the performance metrics of R^2^, MAE, and MSE [mean (standard deviation)] over 5-fold cross-validation) of various regression models for predicting boll length and cellulose content. The results highlight the excellent performance of the Random Forest model in both cases, demonstrating the complementary value of integrating nanomechanical (AFM) and transcriptomic (CESAs) features for cotton fiber trait prediction.

| Target | Model | $R^2$ | MAE | MSE |
| --- | --- | --- | --- | --- |
| Boll Length | LinearRegression | 0.90 (0.03) | 0.24 (0.02) | 0.10 (0.02) |
|  | Ridge | 0.90 (0.04) | 0.23 (0.03) | 0.09 (0.03) |
|  | Lasso | 0.91 (0.04) | 0.22 (0.03) | 0.09 (0.03) |
|  | ElasticNet | 0.91 (0.04) | 0.23 (0.02) | 0.09 (0.03) |
|  | RandomForest | 0.90 (0.05) | 0.22 (0.03) | 0.09 (0.03) |
| Cellulose Content | LinearRegression | 0.79 (0.07) | 0.31 (0.07) | 0.21 (0.09) |
|  | Ridge | 0.80 (0.07) | 0.30 (0.07) | 0.21 (0.09) |
|  | Lasso | 0.78 (0.06) | 0.31 (0.06) | 0.22 (0.08) |
|  | ElasticNet | 0.79 (0.06) | 0.31 (0.05) | 0.21 (0.08) |
|  | RandomForest | 0.94 (0.03) | 0.14 (0.03) | 0.07 (0.04) |

AFM-only models, underscoring the complementary nature of gene expression and nanoscale structural and mechanical features in capturing key determinants of fiber development. Table 1 demonstrates the prediction capability of our two target variables in far-ranging ML prediction models.

We further assess the influence of each feature on the predictions by performing SHAP analyses [32] at three distinct developmental stages (e.g., 6–10 DPA, 12–14 DPA, and 16–20 DPA). Fig. 3 (c-d) show that for boll length, the number of microfibrils crossovers consistently emerges as the top predictor across all stages, followed by microfibrils angle (among top three predictors). This suggests that the structural complexity of microfibrils plays a central role in fiber elongation. Several secondary wall–related CESA genes (e.g., CESA4, CESA7, CESA8) appear among the most influential features (among the top 10), highlighting the importance of cellulose synthase activity as fibers transition from primary to secondary wall formation. However, the persistent placement of CESA6, a primary wall-related CESA, in the top three predictors across all three developmental windows is interesting. In general, CESA6 is known for its role in primary wall synthesis, and most CESA6 paralogs in Gh and Gb decrease as development progresses. Only exception is CESA6B that may increase in expression over development for both homoeologs.

**Fig. 3.**
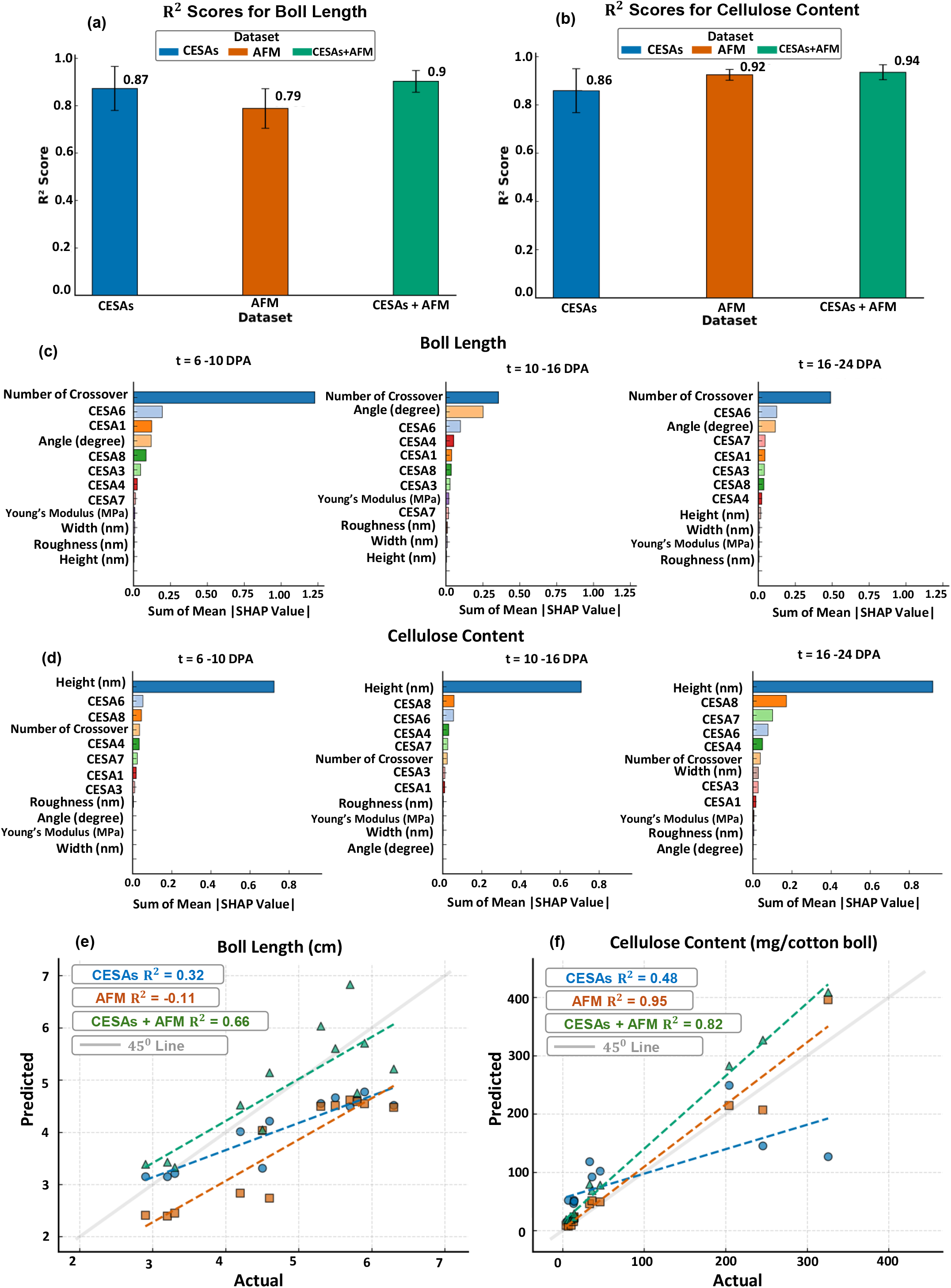
Random Forest model performance (R^2^ across folds) and SHAP-based feature importance for predicting Boll Length and Cellulose Content (a–d). Here, the significance of combining transcriptomic (CESAs) and nanomechanical (AFM) characteristics for cotton fiber trait prediction is illustrated. (a–b) R^2^ scores for CESAs, AFM, and CESAs + AFM datasets across 5 folds where error bars are highlighting the standard deviation. (c–d) Grouped SHAP value summaries for different developmental stages (6–10, 10–16, and 16–24 DPA), highlighting key predictors from CESAs and AFM features. Model predictions on high-quality data: *Gossypium barbadense (12 test data points*) when trained on low-quality data: *Gossypium hirsutum* (e, f). Scatter plots for (e) Boll Length and (f) Cellulose Content compare predicted versus actual values across CESAs, AFM, and CESAs + AFM datasets. Dashed lines indicate best-fit regressions for each dataset, with corresponding R^2^ values annotated.

**Fig. 4.**
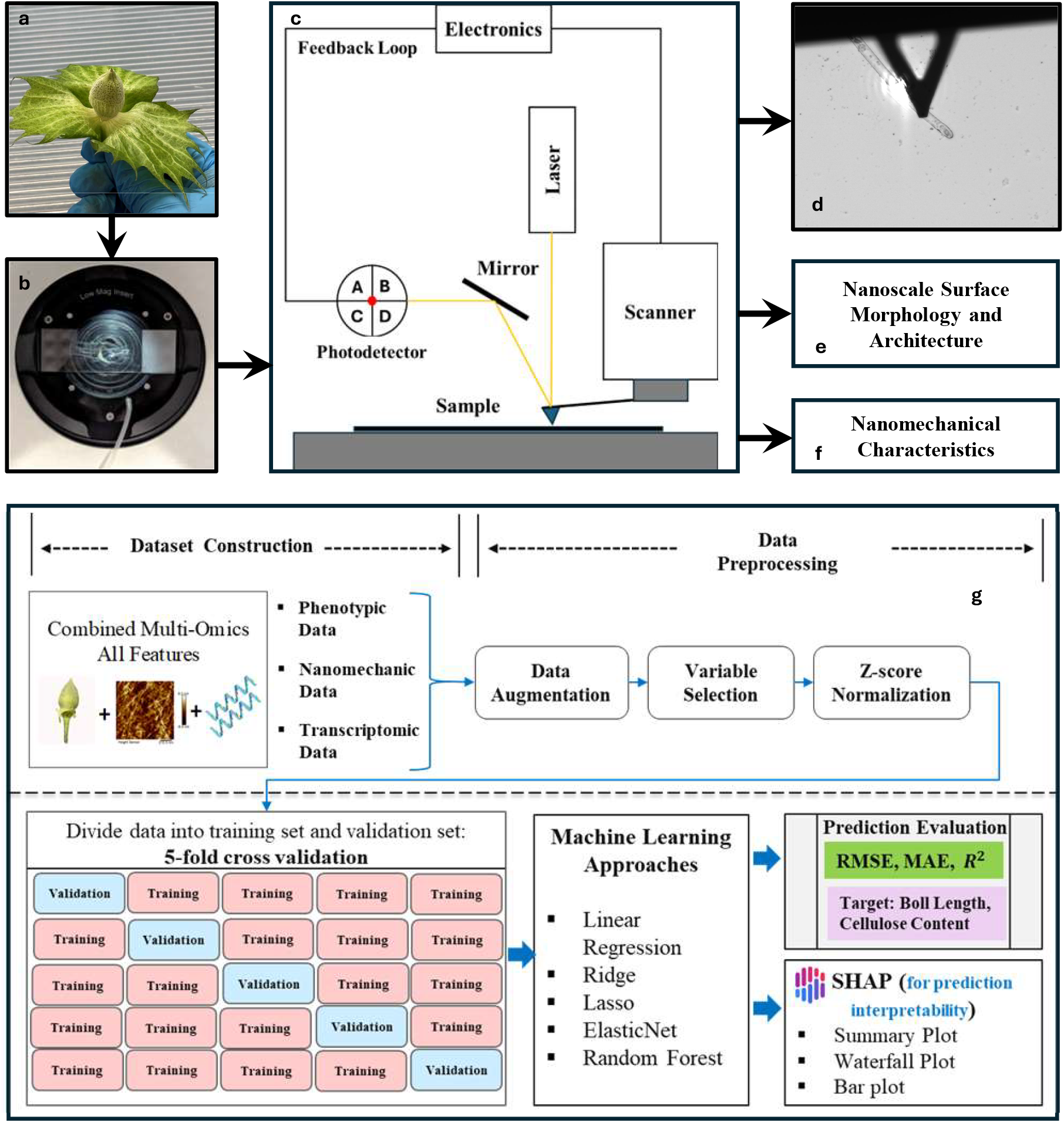
Overall workflow for conducting the study on Cotton Fibers. (a) Cotton boll sample harvested from the cultivated chamber. (b) Fiber sample preparation setup, illustrating how fibers are mounted on the glass slide to ensure stability for AFM nanoscale imaging. (c) Schematic representation of the AFM system, showing the laser, controller, and scanning process for imaging and nanomechanical characterization of microfibrils. (d) Microscopic view of the AFM cantilever tip interacting with the surface of a single fiber to acquire structural and nanomechanical data. (e) Nanoscale surface morphology and architecture captured through AFM imaging. (f) Nanomechanical characterization of fiber microfibrils, providing insights into their Young’s modulus and roughness behavior. (g) Overview of the phenotypic feature prediction workflow, showing each step from data pre-processing through modeling and explainability. First, raw data undergoes cleaning and transformation to create suitable inputs for machine learning. Next, the modeling phase outlines how data are split and which regression models are employed. Finally, model performance is evaluated, and SHAP-based interpretability methods reveal the contribution of individual features to the predicted phenotypes.

In contrast to boll length, cellulose content is consistently associated with microfibrils height, indicating that fiber dimensions strongly correlate with cellulose deposition. This feature is the only one to rank among the top three predictors for any developmental window. Most top-ranked predictors are CESAs, although the rank change during development. During the first time window, the primary wall associated with CESA6 is the second greatest predictor. However, it consistently drops in rank as development progresses. Similarly, the secondary cell wall associated with CESA8 improves in rank nearly immediately, followed by the other secondary cell wall CESA genes, CESA4 and CESA7. Although the expression of all CESA genes ranks higher than most nanoscale parameters with respect to cellulose content, additional AFM parameters (angle and crossover count) are among the top 10 predictors. This confirms that both molecular drivers and fiber nano-structure influence overall cell wall accumulation.

Because the current fiber phenotypes possessed by Gh and Gb reflect their independent domestication histories with potentially different targets of selection, we test the generalizability of our model by training the model on Gh and predicting on Gb. Again, the combined (CESAs+AFM) model yield the best performance (Fig. 3 (e, f)), achieving *R*^2^ = 0.66 for boll length and *R*^2^ = 0.82 for cellulose content. In contrast, AFM-only models show poor generalization (negative *R*^2^ for boll length Fig.3(e)), emphasizing the need to incorporating gene expression data. The negative R^2^ is likely because most of the predicted values fall below the mean of the actual values. The negative R^2^ value observed for boll length in the AFM-only model reflects poor generalization performance, likely due to high phenotypic variation between species and the lack of genetic context. Given the different domestication histories and selection pressures of Gh and Gb, incorporating gene expression data, especially CESAs proves to be essential in improving cross-species predictive accuracy. Taken together, these findings highlight the robust predictive capability of combining CESAs and AFM features across multiple developmental stages and species for understanding and forecasting key cotton fiber traits.

## Discussion

This study presents a novel multi-scale investigation of cotton fiber development, focusing on nanomechanical, transcriptomic, and macroscale phenotypic traits in Gh and Gb, which remain largely unexplored to date. While previous research examined transcriptomics and fiber traits [3, 33, 34], difference between nanoscale structural and mechanical properties of these species have not been systematically characterized. Our findings demonstrate that Gb fibers exhibit better microfibril organization, enhanced Young’s modulus, and lower surface roughness across growth stages (8, 12, 18, and 22 DPA), that can be directly correlated with their higher tensile strength and improved textile quality. Use of AFM in PeakForce QNM air mode helps reveal fundamental differences in nanoscale properties between Gh and Gb. A key distinction between Gh and Gb is the timing and nature of cellulose deposition, which might be influencing fiber strength at both the nanoscale and macroscale levels. As illustrated in Fig. 1(a), cellulose accumulation in Gh begins rapidly after 16 DPA, whereas Gb maintains a more gradual increase until 19 DPA. This extended elongation phase in Gb might help ensure precise microfibril alignment and cellulose deposition, resulting in smoother, stronger, and more elastic fibers. This process is driven by sustained turgor pressure (*P*), as explained by Cosgrove’s biomechanical model: *Gr* =*ϕ*(*P-Y*), where growth rate (*Gr*) depends on turgor pressure, extensibility (*ϕ*), and yield threshold (*Y*). The role of water uptake in cell expansion is further expanded as: 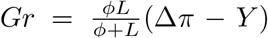. Osmotic pressure difference across the plasma membrane is Δ*π* and the hydraulic conductance is *L* [35]. These models reflect how water enters the growing cell and provide factors into the growth rate due to water transport dynamics, leading to fiber elongation. During that elongation phase, the fibers grow in length while maintaining a consistent diameter, creating a strong and uniform structure [8, 9]. Gb fibers might exhibit prolonged plasmodesmatal closure and enhanced expression of sucrose and K^+^ transporters, maintaining higher turgor pressure (*P*) over extended periods, ensuring sustained elongation. Furthermore, the fiber specific *β*-1,3-glucanase gene facilitates elongation by reopening plasmodesmata, supporting continued nutrient flow and elongation [36], enabling Gb to produce longer fibers than Gh.

During the early elongation phase (8-12 DPA), both Gh and Gb exhibit high crossover densities that indicate the PCW organization. During the transition from PCW to SCW, transcriptionally regulated cell wall-degrading enzymes break down the cotton fiber middle lamella (CFML), releasing fibers as individuals to facilitate SCW formation and altering the orientation of cellulose microfibrils [36, 37]. As a result, by 18 and 22 DPA, both Gh and Gb show a significant reduction in crossover density and a more refined microfibrillar structure (Fig. 1(d-k) and Fig. 2(a-h)). The microfibrils crossover count per 500 × 500 nm^2^ area ( Fig. 2(i, j)) further reveals distinct structural differences during fiber growth, with Gb fibers exhibiting a higher frequency of crossover events. This may reflect increased microfibrils density during cell wall synthesis, suggesting a more compact and cohesive architecture in Gb fibers compared to Gh.

The earlier transition to SCW synthesis in Gh could be a reason for bulkier cotton bolls, less ordered microfibrils, leading to increased surface roughness and reduced mechanical performance. Also, the rapid cellulose deposition correlates with reduced extensibility (*ϕ*), impairing stress relaxation and wall loosening efficiency [38]. At 18 and 22 DPA, Gh fibers exhibit a rougher surface compared to Gb fibers (Fig. 2(k)). This confirms that Gb fibers undergo more precise cellulose microfibril organization and cellulose compaction during SCW deposition, leading to smoother fibers with higher durability, reduced pilling, and improved dye absorption [2]. The nanomechanical measurements (Fig. 2(l)) show that Gb fibers consistently exhibit higher Young’s modulus values compared to Gh due to more organized microfibril arrangement, enhanced hydrogen bonding, and a better cellulose deposition pattern, leading to higher tensile strength and elasticity at the macroscale. The differences in cellulose microfibril dimensions between both species also align with their insights on elongation, cellulose deposition, PCW to SCW transition, and bulkiness of cotton bolls. At 8 and 12 DPA, these dimensions for both species are minimal, corresponding to the elongation phase and PCW formation. Cellulose deposition increases drastically just after the end of the elongation phase. Meanwhile, cell wall loosening dominates this elongation phase, enabling stress relaxation and water uptake to drive cell expansion [38]. The transition from PCW to SCW deposition in both Gh and Gb (16 and 19 DPA, respectively) shows a significant developmental change. By 18 to 19 DPA, SCW synthesis leads to a sharp increase in microfibril diameter and height due to a significant amount of cellulose deposition. At 22 DPA, these dimensions increased further. Fig. 2(m, n) demonstrates the comparative analysis of microfibril dimensions. The smaller diameter and height in Gb can be attributed to its prolonged elongation phase, continued wall loosening, and slower cellulose deposition, resulting in compact structures. The tightly packed arrangement of microfibrils can also be a factor in reducing thickness (diameter and height) while maximizing the fiber strength and flexibility. In contrast, Gh fibers have faster cellulose deposition (Fig. 1(c)), leading to bulkier microfibrils with less precise alignment of the individual microfibrils with each other. The compact microfibrils of Gb balance elongation and deposition, whereas Gh prioritizes rapid growth and yield at the expense of refinement and quality. However, tight binding of matrix polysaccharides to cellulosic microfibrils and amorphous glucan chains may increase the microfiber bundle’s apparent diameter and thickness, potentially leading to an overestimation of physical properties measured by AFM or other techniques [39, 40]. Moreover, the overall trend of increasing dimensions remains consistent, ensuring that these variations do not impact the validity of our hypothesis. A more detailed explanation of polysaccharides impacting the thickness of microfibrils is in supplementary section 1.4.

Additionally, the results of our SHAP analysis [32] (Fig. 3) reflect different aspects of cotton fiber development. Some nano-mechanical (AFM) features show a strong association with phenotypic traits such as boll length, which may attribute the fiber length. We find that among these, the microfibrils crossover count during development is the most predictive, perhaps due to its influence on fiber extensibility and strength. In contrast, CESAs gene expression tends to exhibit a more complex relationship with measured phenotypes despite the crucial role of CESAs genes in fiber development. This complex relationship results in lower individual effects of CESAs genes in the prediction models. Moreover, the observed correlation between the transcriptomic and nanomechanical features suggests that there may be underlying interactions that a standard SHAP analysis might not fully capture. Future analyses, such as a causal SHAP approach, may provide additional insight into the interplay among the factors surveyed here. This observation is reiterated in those models based solely on AFM features that exhibited lower predictive quality, with occasional negative *R*^2^ values. These observations underscore the complexity of fiber development and the limitations of relying on a single data type.

An appropriate comparison is presented in Table 2, where existing previous researches are systematically documented to represent the difference between the present research. In our case, we use multi-omics data and exploit machine-learning models for prediction. Additionally, several in-depth analyses are conducted, including single-omics comparative performance. Our investigation includes appropriate feature set findings that can effectively work for phenotypic feature prediction. Fig. 3 illustrates that both target multi-omics datasets perform magnificently. Also, it is noticed that among five different models, which model provides the best prediction, and for each case, how random forest is performing effectively is set down in supplementary Tables 5,6 (supplementary section 1.6). Furthermore, we demonstrate how our experimental design leads to improved predictive performance, confirming the effectiveness of our approach. The improved performance achieved by integrating both the transcriptomic (CESAs) and AFM datasets demonstrates that combining molecular and nano-mechanical data not only enhances prediction accuracy but also provides a more comprehensive understanding of the biological processes at play. Overall, these findings suggest that expanding the dataset and incorporating additional features from both domains could further improve model robustness and accuracy, ultimately supporting more effective cotton breeding strategies and fiber quality enhancement.

**Table 2.** A Comparative Overview of Existing Single/Multi-omics Data and Machine Learning Approaches with the Proposed Methodology.

| Reference and Year | Problem Type | Data Type | Omics Type | Machine Learning Approach | Key Findings |
| --- | --- | --- | --- | --- | --- |
| [41] (2022) | Prediction | Numerical (synthetic and real-world datasets) | Phenotype, genomic | Elastic Net, LASSO, SVR, XGB, MLP | Analyzed prediction using synthetic data and concluded that "Bayes B" outperformed all models in 29/36 cases. |
| [42] (2019) | Prediction (Seed Yield or SY Prediction). | Numerical data (phenotypic) | Phenomics | Random Forest (RF), Recursive Feature Elimination (RFE), 10-fold cross-validation, z-score normalization for predictor variables | Under various agro-management systems SY and phenotypic characteristics are discussed, determined crucial predictor factors, average increase of 56.8 kg/ha in IA-RS; Mean SY deviation in IA-RS was $59 \pm 115$ kg/ha. |
| [43] (2022) | Prediction (plant breeding in general, mentioned maize and wheat) | Numerical, categorical (Multidimensional data including: Genomic, Phenotypic, Environmental data) | Genotype (G), environment (E), and genotype by environment interaction (GEI) | Machine Learning, Deep Learning and Big Data | 1) Genomic prediction using multi-omics information, big data technology, and AI; 2) explored spatiotemporal models; 3) translated smart breeding into genetic gain. |
| [44] (2018) | Classification problem (identifying DEGs as upregulated or downregulated) | Numerical (RNA-seq) | Transcriptomics | Logistic Regression, Classification Via Regression, Random Forest, LMT, Random Subspace | Discovered 23 most useful features, concluded InfoGain + Logistic Regression system provided the most robust and sensitive DEG prediction, validated by qRT-PCR. |
| [45] (2024) | Prediction (plant disease severity (classes) using gene expression profiles) | Numerical (RNA-Seq) | Transcriptomic (both plant + pathogen transcriptomes) | Support Vector Machine (SVM), linear SVM, Random Forest (RF), XGBoost, and Deep Neural Network (DNN) | Experiment resulted in the discovery of genes not previously associated with plant immunity or pathogen response, explored a variety of feature selecting methods. |
| [46] (2024) | Prediction (Genes prediction which are involved in biosynthesis) | Numerical and text data | Multi-omics (genomics, transcriptomics, epigenomics, proteomics data) | Random Forests, LightGBM, CatBoost, ExtraTrees, XGBoost, Neural Networks | Constructed GS dataset, effectively detected key features for model and used AutoGluon-Tabular algorithm which provided more generalized model for multiple species. |
| [19] (2024) | Prediction (of 6 complex traits) | Numerical | Multi-omics data (genomic (G) + transcriptomic (T) + methylomic (M)) | Ridge regression Best Linear Unbiased Prediction (rrBLUP) and Random Forest (RF) | Compared similarity matrices of omics, compared the benchmark genes with the essential genes, investigations are conducted into feature interactions and models that combined multi-omics data performed optimally. |
| [47] (2023) | Prediction (of cotton fibers' 4 main quality indicators) | Time-series data (from UAV (Unmanned Aerial Vehicle)) and RGB image (remote sensing) | Remote sensing data | Deep learning algorithm for image segmentation, machine learning model (neural network) for prediction | Neural network model improved their data prediction accuracy compared to linear regression. Also, model predicts cotton fibers' average length, uniformity index, and micronaire value. |
| Proposed research work | Prediction (cotton fiber boll length and cellulose content) | Numerical | Multi-omics data (phenotypic + nano-mechanical + transcriptomic) | Data preprocessing, Linear regression, Ridge regression, Lasso regression, ElasticNet, and Random Forest (RF) | Various feature analyses are done for prediction performance; it is found that Random Forest performs rigorous growth stage-wise SHAP analysis for explainability. |

## Methods

Overall workflow for this present investigation ( Fig. 4) illustrates how the experimental and computational outputs are derived from cotton bolls through AFM nanoscale analysis. The workflow integrates controlled growth, precise sample preparation, advanced AFM techniques, and machine learning (ML) integration (with multi-omics data) to analyze the structural and nano-mechanical properties of cotton fibers. The following subsections outline the key steps in the experimental process, starting with the environmental and growth conditions for cultivating cotton plants. This also includes sample preparation for AFM experiments to preserve cellulose microfibril structures for nanoscale analysis. The configuration and operational parameters of the AFM setup are described, followed by the methods used for data analysis and characterization, and transcriptomic data generation. Finally, these are integrated with the ML models.

### Environmental and Growth conditions

This study focuses on two prominent commercially cultivated species of cotton plants: *Gossypium hirsutum* (Gh) and *Gossypium barbadense* (Gb). Plant samples were grown in a controlled environment chamber (Conviron E15, Controlled Environments, Inc., Pembina, ND, USA) under carefully regulated conditions. Each plant was individually cultivated in a two-gallon pot containing a potting mix with a 4:2:2:1 ratio of soil, perlite, bark, and chicken grit. The chamber was set to a 16-hour photoperiod with a light intensity of 500 µmol and maintained at a constant temperature of 28°C. Flowers were self-pollinated (anthesis), tagged and the day of anthesis was recorded as 0 DPA. Bolls from 8, 12, 18, and 22 DPA were collected at mid-day, labeled, and stored at −80°C for the subsequent experimental data collection.

### AFM sample preparation

Preparation of cotton fiber samples for Atomic Force Microscopy (AFM) followed a structured approach to achieve the best imaging and characterization results. In this experiment, we considered four consecutive critical days in the growth stages of cotton bolls as 8, 12, 18 and 22 DPA. Cotton bolls, stored at −80°C, were first brought to room temperature to restore their natural state. The boll wall of the cotton bolls was carefully removed using fine-tipped tweezers (Cole-Parmer, Straight Fine Tips, Serrated, 101mm, UX-07398-22) to extract the fibers inside without causing damage and minimizing mechanical stress. Then the fibers were incubated in 0.5M Potassium Hydroxide (KOH) solution at room temperature for 30 minutes to remove surface contaminants. Afterwards, they were rinsed gently in deionized (DI) water for 3 minutes to neutralize and clean the surface. The fibers were then positioned on AFM-compatible Fisherbrand Double Frosted Microscope Slides (Pre-cleaned, 25×75×1.0 mm, Cat. No. 22-034-486) and allowed to air-dry at room temperature for 10 minutes to ensure proper adhesion and precise high-resolution AFM imaging.

### AFM setup for the experiment

The AFM experiment was performed using the BioScope Resolve system (Bruker), which is optimized for detailed and high-resolution analysis.

A Bruker SNL-10 probe (Sharp Nitride Lever) was used for the AFM experiment that combines a sharp silicon tip with a sensitive silicon nitride cantilever. The probe had a tip radius of 2 nm, a spring constant of 0.35 N/m, and a tip height ranging from 2.5 to 8.0 µm. Its triangular cantilever geometry provided stability during operation, with a cantilever thickness of 0.6 µm. A reflective gold coating on the back side of the cantilever enhanced laser detection and ensured precise sample interaction. AFM imaging was performed in PeakForce QNM air mode inside a vibration and noise isolation chamber to minimize external noise. Before performing each experiment, the deflection sensitivity and cantilever spring constant were calibrated to maintain measurement accuracy. The imaging parameters were set to a scan rate of 0.450 Hz, a peak force setpoint of 750 pN, and a peak force amplitude of 250 nm. High-resolution scans were conducted with a resolution of 256 × 256 pixels per line, and the peak force frequency was consistently maintained at 1 kHz throughout the experiments to obtain the nanoscale topographical and mechanical properties of the cotton fibers.

### Data analysis and characterization

For detailed analysis of the nanoscale structural and mechanical properties of cotton fiber microfibrils across different growth stages (8, 12, 18, and 22 DPA), multiple software tools and statistical methods were employed to ensure accurate and comprehensive evaluation. Gwyddion software was used to measure the diameter, height, and surface roughness of the microfibrils. NanoScope Analysis software was utilized to calculate Young’s modulus (by fitting force-distance curves to the Hertz model, as described and outlined by Masud et al. [25] and supplementary section: 1.2. Young’s modulus determines the mechanical strength of the microfibrils. For each developmental stage, 50 randomly selected single microfibrils were analyzed, and average values were calculated to ensure consistency and reliability. To further examine the microfibril architecture, MATLAB image processing (details available in supplementary section: 1.3 and Supplementary Fig. S4, S5) was employed to measure angles between microfibrils, determine crossover counts, and analyze correlations and intensity variations within the microfibrils. The average values were calculated to present the representative data. Finally, a normal t-test was conducted to evaluate the statistical significance of the measured properties, ensuring that the observed differences were meaningful enough to interpret. The macroscale structural and physiological properties of cotton fibers were assessed using a combination of direct measurements and previously reported data to provide a comprehensive comparison between Gh and Gb. The cotton boll length for Gb was measured using a digital slide caliper (Fisherbrand Traceable Digital Calipers, resolution: 0.01mm, Catalog No.14-648-17), which recorded the distance from the tip to the base of the boll. For Gh, the boll length data were extracted from the study reported by Wilson et al.[33]. The crystalline cellulose content of the fibers for Gh and Gb were obtained from our collaborator’s study and it was estimated by Updegraff reagent[3].

### Transcriptomic data generation for Gb and Gh

For the RNA transcriptomics, previously generated RNA-seq data from developing cotton fibers (6-25 days DPA) growth under controlled conditions was downloaded from NCBI (PRJNA1099209) [34]. Reads were quality-trimmed using Trimmomatic (v0.39) and pseudoaligned to a species-specific, homoeolog-diagnostic G. raimondii reference transcriptome using Kallisto (v0.46.1). Gh and Gb homologs were identified by mapping the transformed transcriptome to each genome using gmap [48, 49] and intersecting the annotation locations using bedtools2 [50]. In the present study, we focused on the expression of cellulose synthase genes (CESA), which play important roles in fiber development. In both Gb and Gh (supplementary Tables: 1,2,3,4), most PCW CESAs homologs (i.e., CESA1/3/6) exhibit decreasing expression throughout the developmental time course, as expected from their diminishing roles as the fiber progresses into SCW synthesis. Conversely, the SCW CESAs genes (i.e., CESA4/7/8) generally exhibit increasing expression as the fiber progresses through development and transitions toward secondary wall synthesis. Notably, two exceptions exist. In both Gh and Gb, both homoeologs of the PCW CESA6 paralog CESA6B increase in expression across development whereas the SCW CES8A paralog decreases in expression as the fiber moves into SCW synthesis. The significance of the contradictory expression found in these two CESA paralogs is yet unknown and warrants further exploration.

### Multi-omics Data and Machine Learning Integration

#### Dataset Preparation

For machine learning prediction, we assembled a comprehensive dataset that integrates phenotypic, transcriptomic, and nano-mechanical features from both high-quality (Gb) and low-quality (Gh) cotton fibers. The phenotypic data, obtained from [33], comprise measurements of fiber boll length (cm) and cellulose content (mg per cotton boll). The transcriptomic dataset includes 29 TPM values corresponding to cellulose synthase (CESA) genes, derived from processed RNA-seq data. These measurements were recorded daily from 5 to 25 DPA, with three replicates per time point [33]. Additionally, nanomechanical properties—computed as described in the previous section include microfibril width (nm), height (nm), surface roughness (nm), Young’s modulus (MPa), angular orientation (°), and the number of crossovers.

#### Data Augmentation

Collecting large amounts of multi-omics data is challenging, so our dataset has varying amounts of data across the different omics types. To reduce bias in our machine learning predictions, we used data augmentation to boost the less-represented features. Specifically, bootstrapping combined with XGBoost was applied to impute missing values, and the data were organized according to the cotton plant growth stages. Including the augmented data, we had 200 data points for low-quality and 12 for high-quality cotton.

#### Variable Selection

Variable selection is essential for dimensionality reduction, computational efficiency, and model interpretability. We used a Gaussian Process Regressor (GPR) visualization technique (Supplementary Fig. S6) to assess feature trends and signal quality, discarding those without clear patterns. Additionally, a correlation matrix was computed to identify and remove transcriptomic features with correlations greater than 97% to reduce multicollinearity. Detailed heatmaps (Supplementary Fig. S7, S8) and GPR plots are provided in supplementary section: 1.5).

#### Z-Score Normalization

Z-score normalization (standardization) transforms numeric data to a mean of 0 and a standard deviation of 1. This harmonizes features measured on different scales, ensuring that all contribute equally to the analysis.

#### ML Models

We evaluated five predictive models: linear regression, ridge regression, lasso regression, elastic net, and random forest regressor. Linear regression models the relationship between the dependent variable *y* and the predictors *X* as [41]

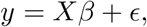

minimizing the mean squared error. Ridge regression extends this framework by adding an L2 penalty, 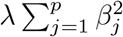, to shrink coefficient magni-regression uses an L1 penalty, 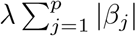wihtch some tudes and reduce overfitting. In contrast, lasso coefficients to zero, promoting feature selection. Elastic net combines both penalties, balancing the benefits of ridge and lasso regularization. In addition, we employed the random forest regressor—a nonlinear ensemble method that constructs multiple decision trees on bootstrapped data subsets and aggregates their predictions—to capture complex feature interactions.

#### Training and Evaluation

To ensure robust performance, we used 5-fold cross-validation along with grid search for hyper-parameter optimization. For ridge and lasso regression, the regularization parameter *α* was tuned over {10^*−*3^, 10^*−*2^, 10^*−*1^, 10^0^, 10^1^}. The elastic net model was optimized over *λ* ∈ {10^*−*3^, 10^*−*2^, 10^*−*1^, 10^0^} and L1 ratios {0.2, 0.5, 0.8}. For the random forest regressor, model complexity was controlled by testing 50, 100, and 200 estimators. Model performance was assessed using Root Mean Squared Error (RMSE),

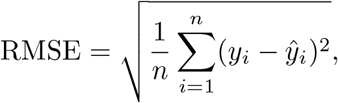

Mean Absolute Error (MAE),

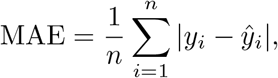

and the coefficient of determination (*R*^2^),

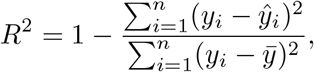

where *y*_*i*_ denotes the true value, *ŷ*_*i*_ referes to the predicted value, and 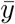 represents the mean of the observed values [51]. The optimal model was chosen based on the highest *R*^2^ and the lowest error metrics.

### Explainability Analysis Using SHAP

We applied SHAP (SHapley Additive exPlanations) to elucidate the contributions of individual features to our model predictions. Based on Shapley values from cooperative game theory, the SHAP value for feature *j* is computed as [52]

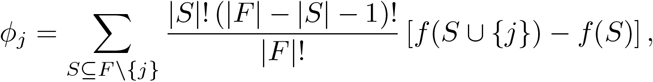

where *F* denotes the complete set of features, *S* is any subset of features excluding *j*, and the weighting factor ensures a fair distribution of contribution.

### SHAP Visualization and Grouping

To capture both global trends and instance-level details, we generated SHAP plots that illustrate feature importance. In our transcriptomic dataset, which includes 29 features related to cellulose synthase (CESA) genes, we grouped features based on their association with specific CESA proteins (1, 3, 4, 6, 7, and 8), as described in the Transcriptomic Data Generation section. Moreover, we organized the SHAP values into three temporal windows—6–10, 10–16, and 16–24 DPA —to better reflect the dynamics of gene expression over developmental stages.

## Consent for publication

Not applicable

## Ethics approval and consent to participate

Not applicable

## Data availability

The entire data set will be available upon request to the corresponding author.

## Materials availability

Not applicable

## Code availability

All the codes will be available upon request to the corresponding author.

## Author contributions

Conceptualization, A.S.; Methodology, A.S., N.M., M.H.H.H, S.S.; Software, A.B., T.Z.J., S.S; Formal Analysis, M.H.H.H, B.A.; Investigation, M.H.H.H, N.M., A.B, T.Z.J. and A.S.; Resources, A.S.; Writing - Original Draft, M.H.H.H, N.M., A.B, T.Z.J., C.G., and A.S.; Writing - Review and Editing, A.S,; Visualization, A.S.; Supervision, A.S., O.A.Z., C.G., J.F.W.; Project Administration, A.S.; Funding Acquisition, A.S., O.A.Z., J.F.W.

## Acknowledgments

We would like to thank Iowa State University for their continued support.

## 1. Supplementary information

### 1.1. Supplementary figures and tables

**Figure S1.**
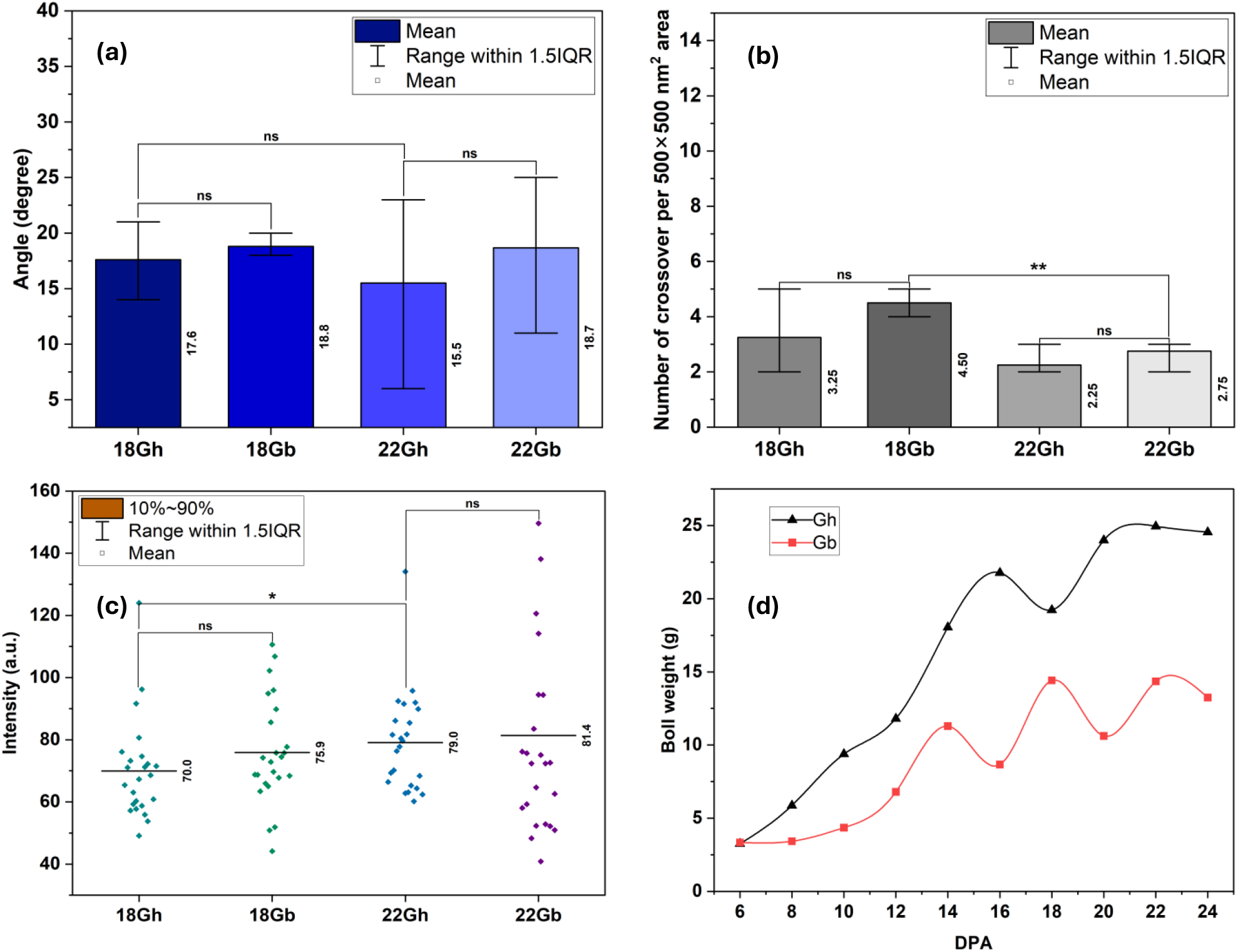
Comparative analysis of microfibril structural properties and boll weight in Gossypium hirsutum (Gh) and Gossypium barbadense (Gb) during fiber development. (a) Microfibril crossover angles for Gh and Gb at 18 and 22 DPA, showing no significant differences between the two species at both stages, as most of the microfibrils become almost straight and mature. (b) Number of microfibril crossovers per 500 × 500 nm^2^ area, exhibiting negligible differences between Gh and Gb. Due to primary to secondary cell wall transformation, it shows very few crossovers of microfibrils at higher DPA. (c) Intensity measurements of microfibrils at 18 and 22 DPA, where Gh and Gb display comparable intensities, except for a minor but significant difference during cell wall transformation (18 DPA to 22 DPA). (d) Boll weight progression from 6 to 24 DPA, showing that Gh consistently develops bulkier bolls than Gb, with an upward trend in weight as DPA increases. Statistical significance is denoted as follows: ns indicates no significance, * represents p *<* 0.05, ** represents p *<* 0.01, *** represents p *<* 0.001, and **** corresponds to p *<* 0.0001.

**Figure S2.**
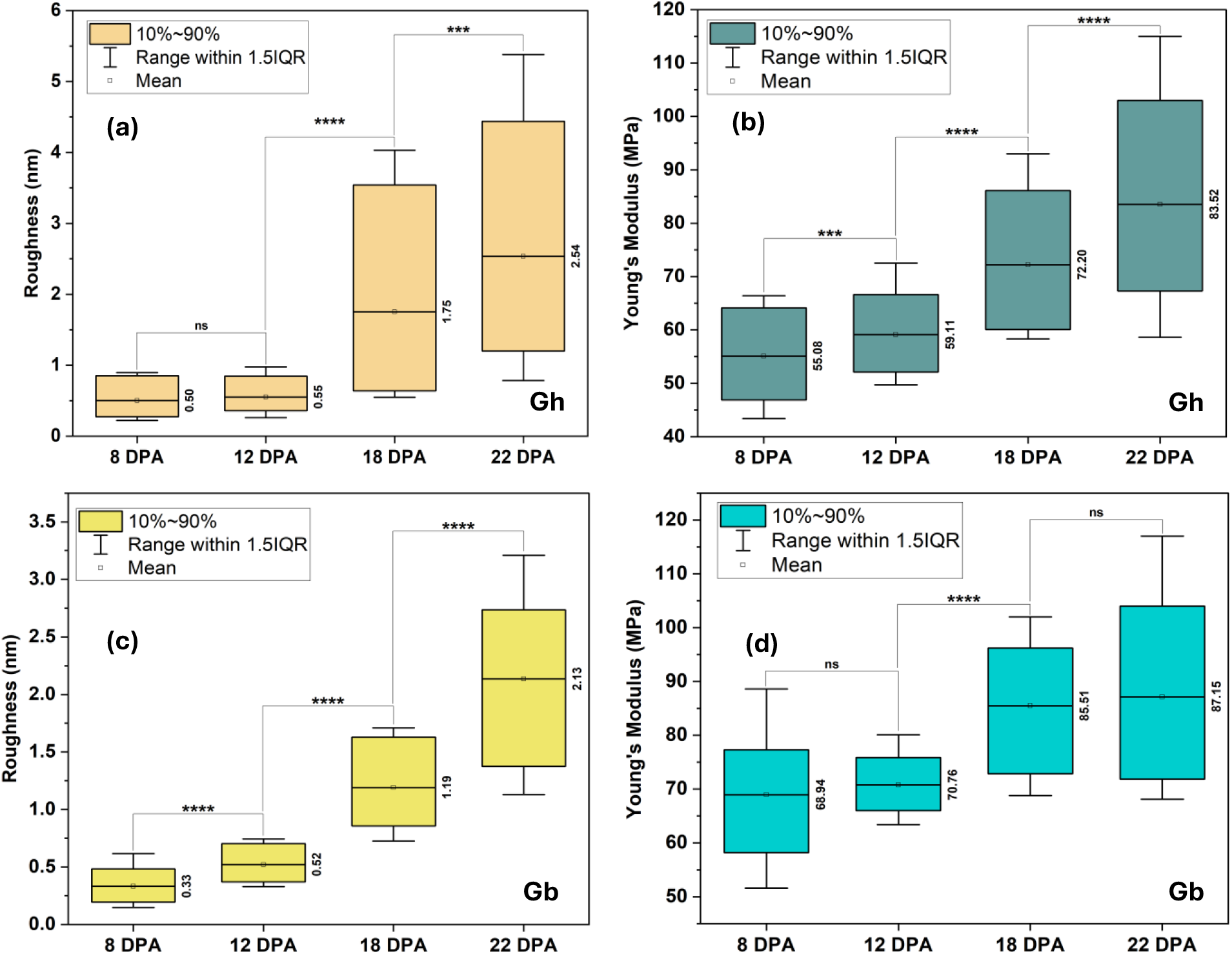
Mechanical properties of microfibrils for Gossypium hirsutum (Gh) and Gossypium barbadense (Gb) across the developmental stages. (a, b) Surface roughness and Young’s modulus of Gh microfibrils progressively increase from 8 to 22 DPA, reaching their highest values at 22 DPA. (c, d) Gb exhibits a similar upward trend but maintains consistently lower roughness and higher Young’s modulus, indicative of its more refined microfibril properties. Statistical significance is denoted as follows: ns indicates no significance, * represents p *<* 0.05, ** represents p *<* 0.01, *** represents p *<* 0.001, and **** corresponds to p *<* 0.0001.

**Figure S3.**
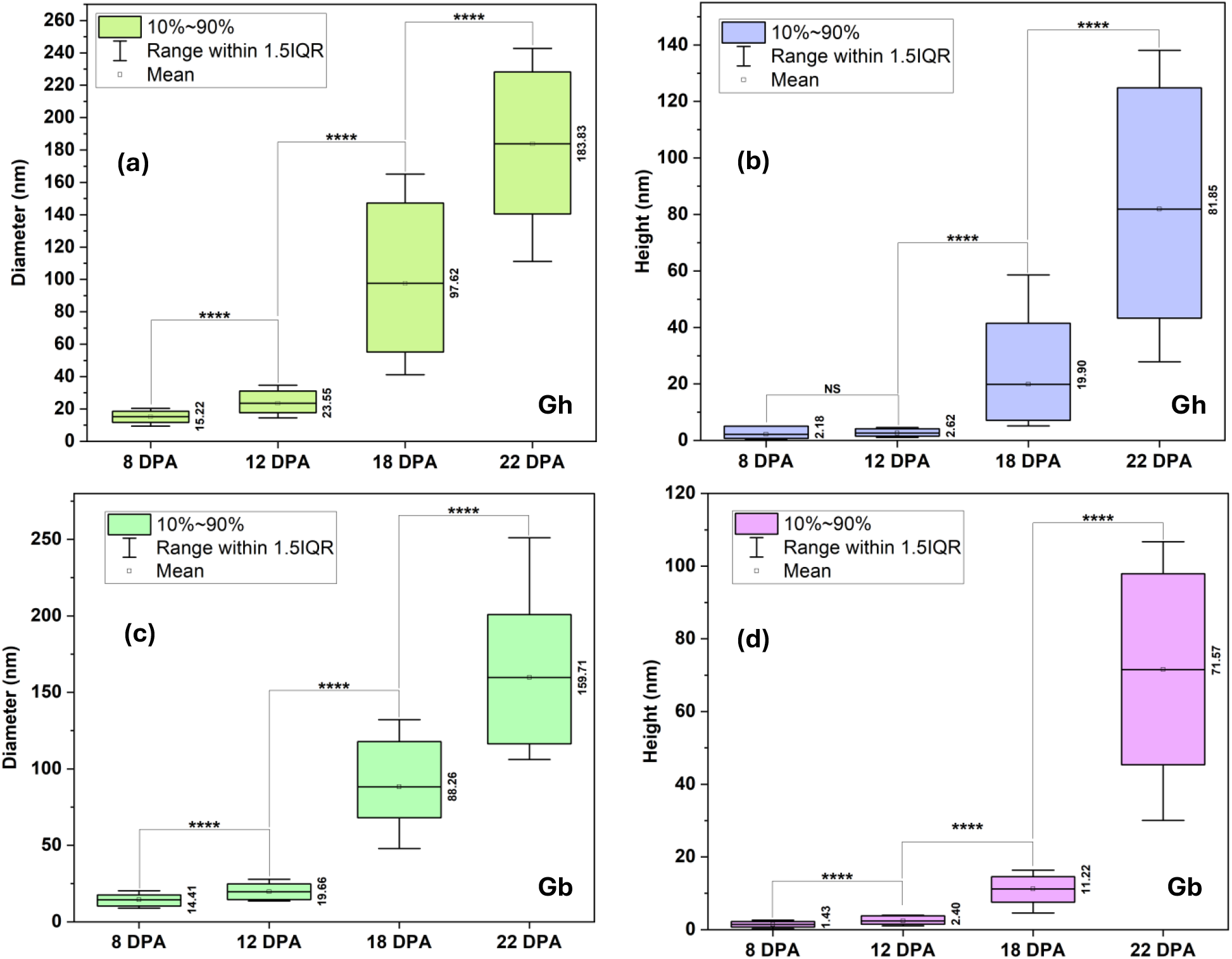
Microfibril dimensions of Gossypium hirsutum (Gh) and Gossypium barbadense (Gb) across the developmental stages. (a, b) Microfibril diameter and height of Gh from 8 to 22 DPA show significant increases, with the largest dimensions observed at 22 DPA. (c, d) Microfibril diameter and height of Gb follow a similar trend but remain consistently smaller, reflecting its finer and more compact structure. Statistical significance is denoted as follows: ns indicates no significance, * represents p *<* 0.05, ** represents p *<* 0.01, *** represents p *<* 0.001, and **** corresponds to p *<* 0.0001.

**Figure S4.**
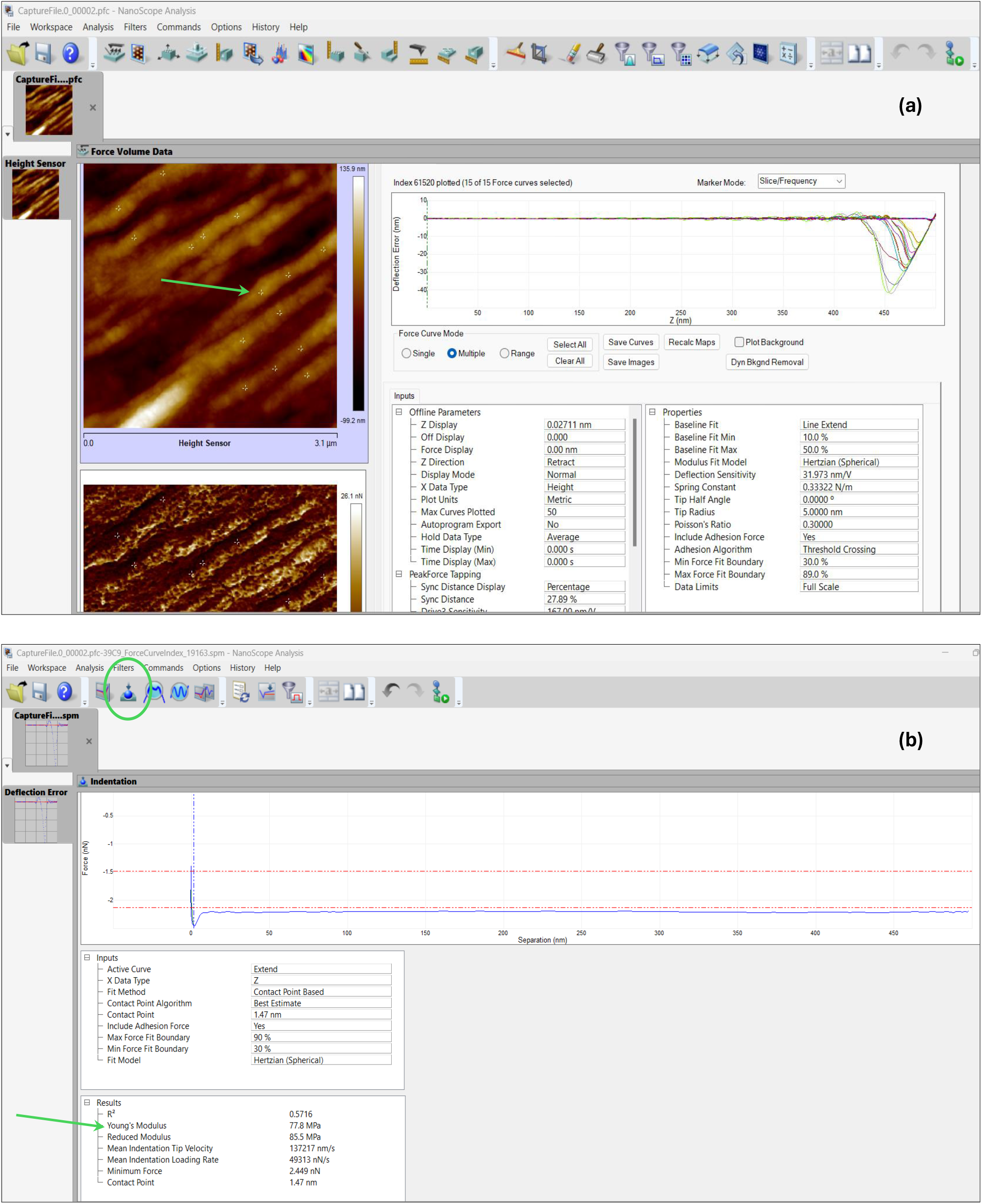
Calculation of Young’s modulus using NanoScope Analysis software from AFM force–distance data. The analysis uses .pfc files obtained during PeakForce QNM AFM experiments. (a) Multiple points are selected along the microfibrils to extract corresponding force–distance curves. (b) The extracted curves are analyzed using the Indentation function in NanoScope Analysis, where Hertzian model is applied to calculate Young’s modulus.

**Figure S5.**
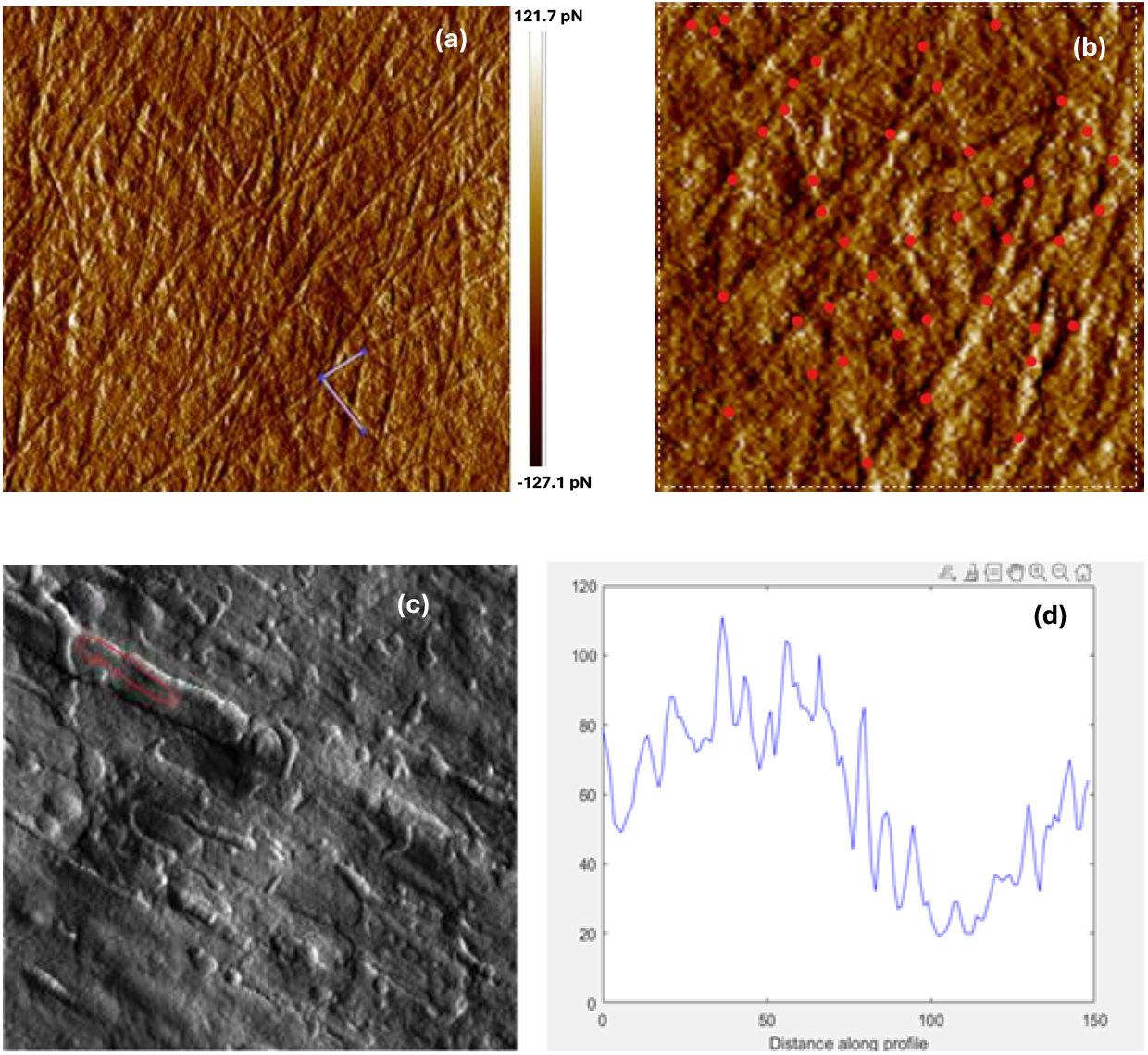
(a) AFM image of Gossypium hirsutum fiber (Gh) at 8 DPA with overlap lines indicating the microfibrils overlap angle of 79 degrees. (b) Microfibrils crossover count analysis for a 8 DPA Gh fiber showing the number of crossovers identified within a 500 nm × 500 nm region. (c) AFM image of Gh fiber at 18 DPA highlighting structural features for intensity measurements. (d) Intensity profile measured along the red-circled region in (c), illustrating intensity variations along the microfibril structure.

**Figure S6.**
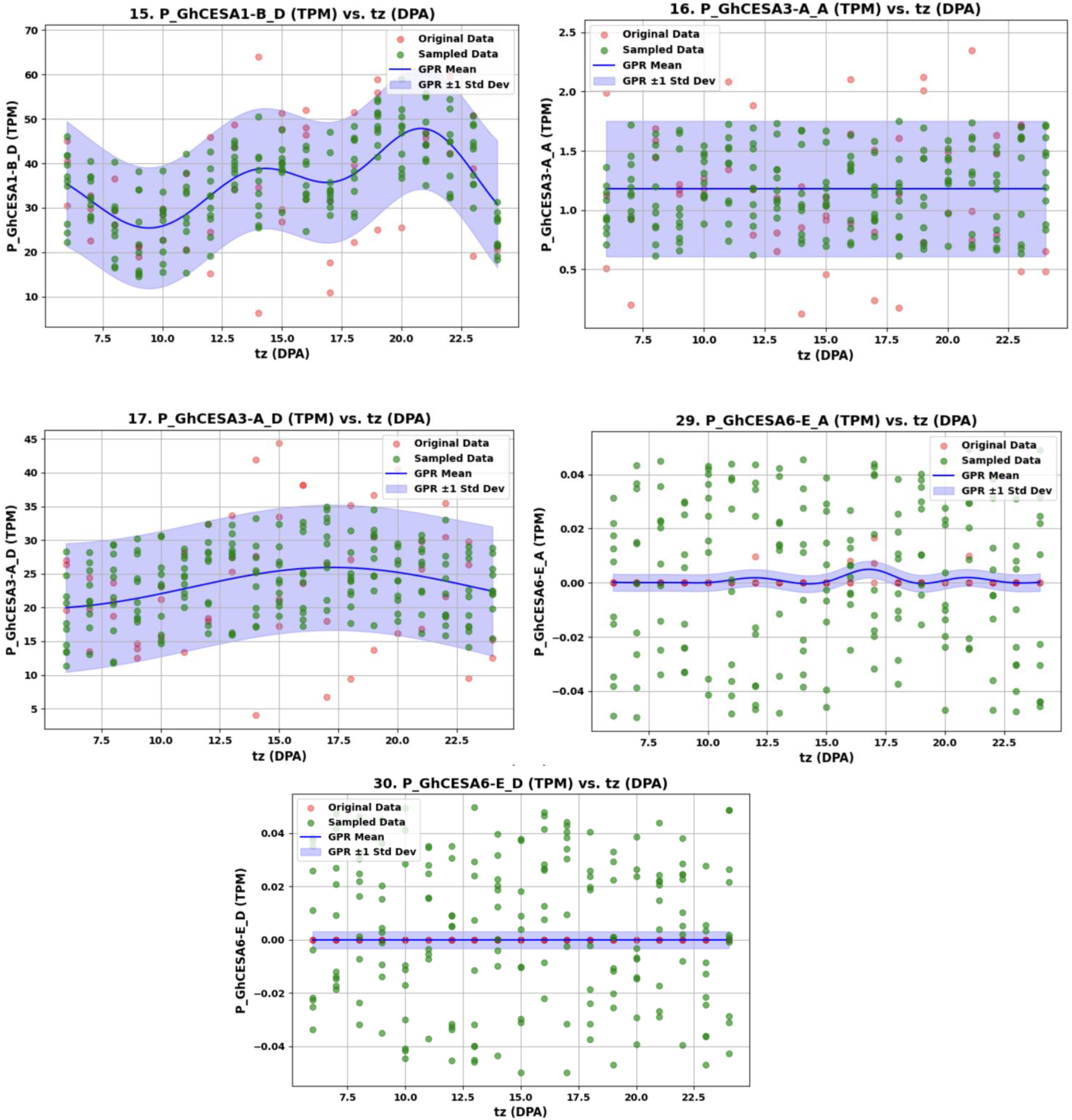
GPR visualization of transcriptomic features. Features that do not exhibit distinct, informative patterns are excluded from subsequent analyses to avoid introducing noise into the ML models.

**Figure S7.**
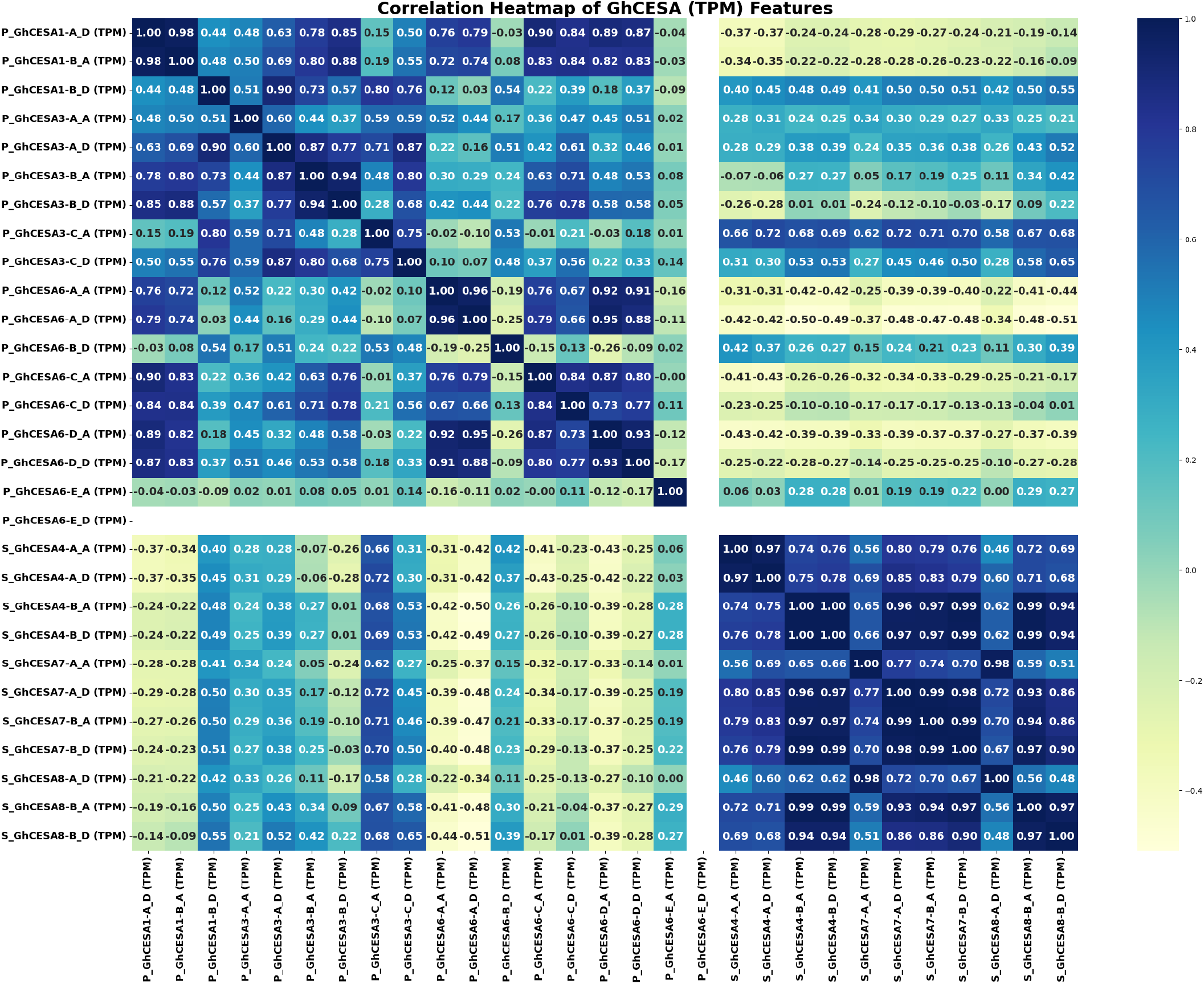
Initial heatmap displaying pairwise Pearson correlation coefficients among transcriptomic features related to cotton fiber development. Dark blue indicates strong positive correlations, while dark red indicates strong negative correlations, revealing clusters of high correlation.

**Figure S8.**
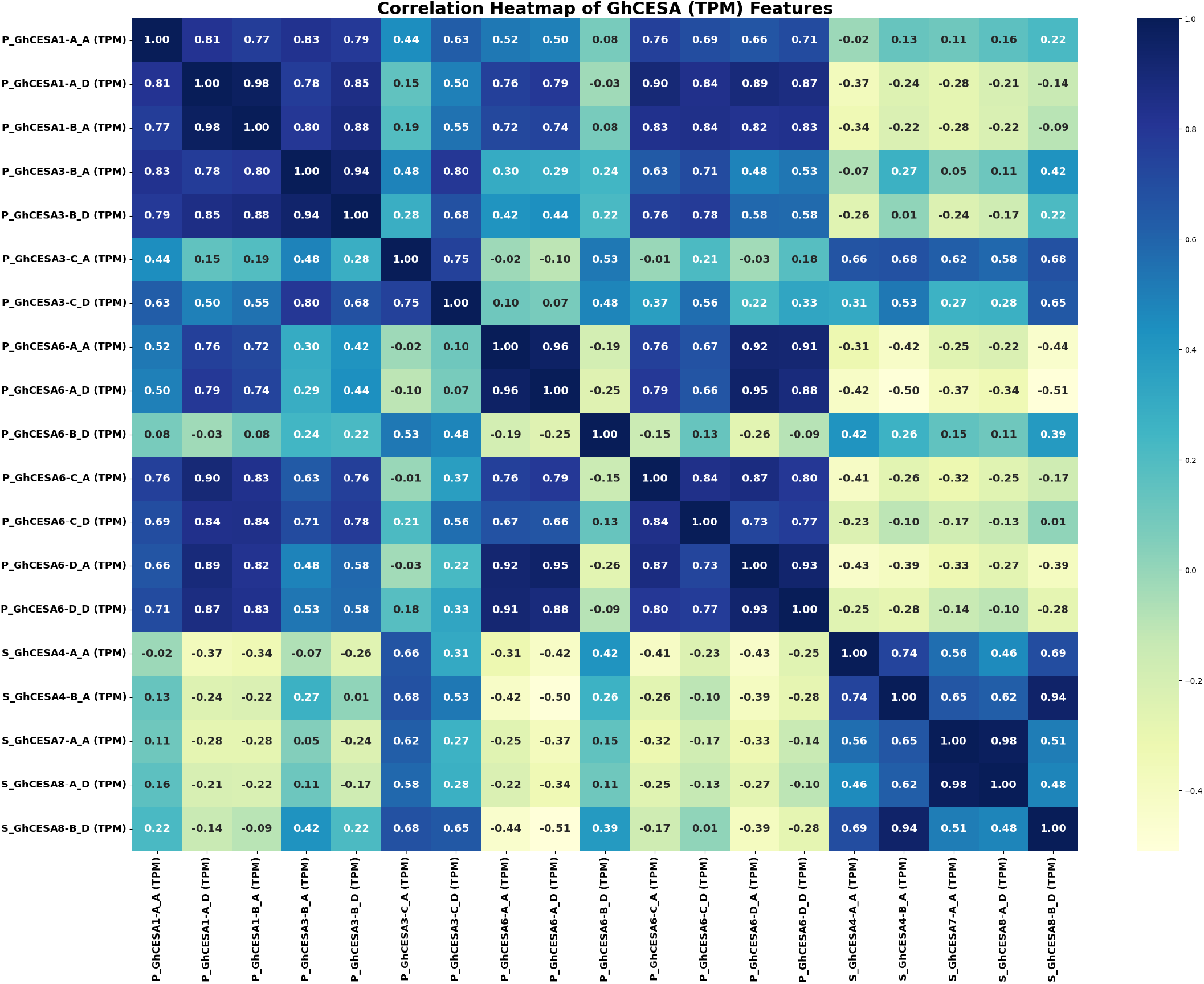
Reduced correlation matrix after applying a threshold of r *>* 0.97. Redundant features have been removed, retaining one representative gene per cluster to maintain biological signals and reduce noise.

**Table 1.**
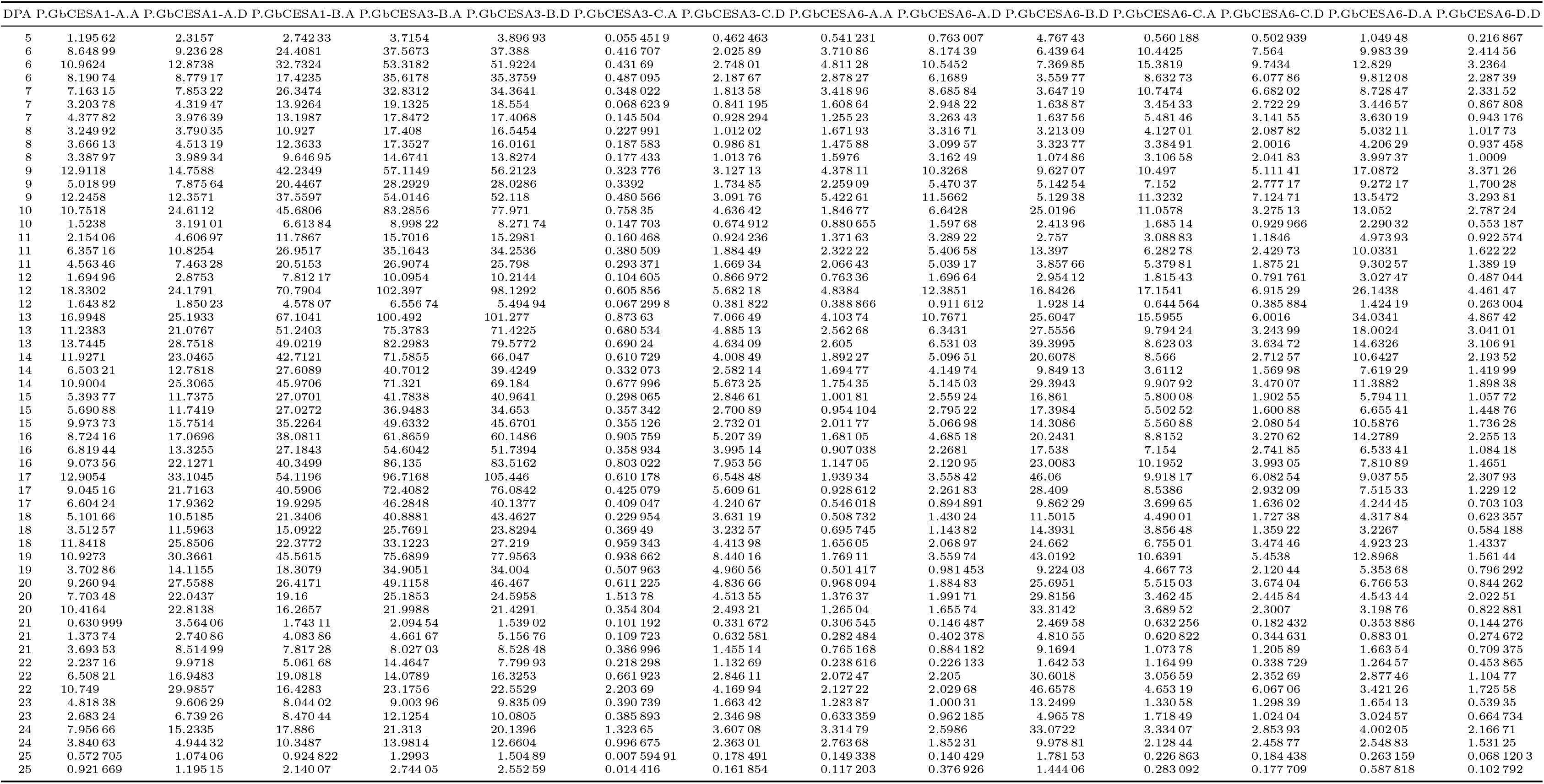
Transcriptomic data of *Gossypium barbadense* (Gb) in Transcripts Per Million (TPM) related to primary cell wall cellulose synthesis. This dataset emphasizes the significant contributions of CESA1, CESA3, and CESA6 in cellulose biosynthesis.

**Table 2.**
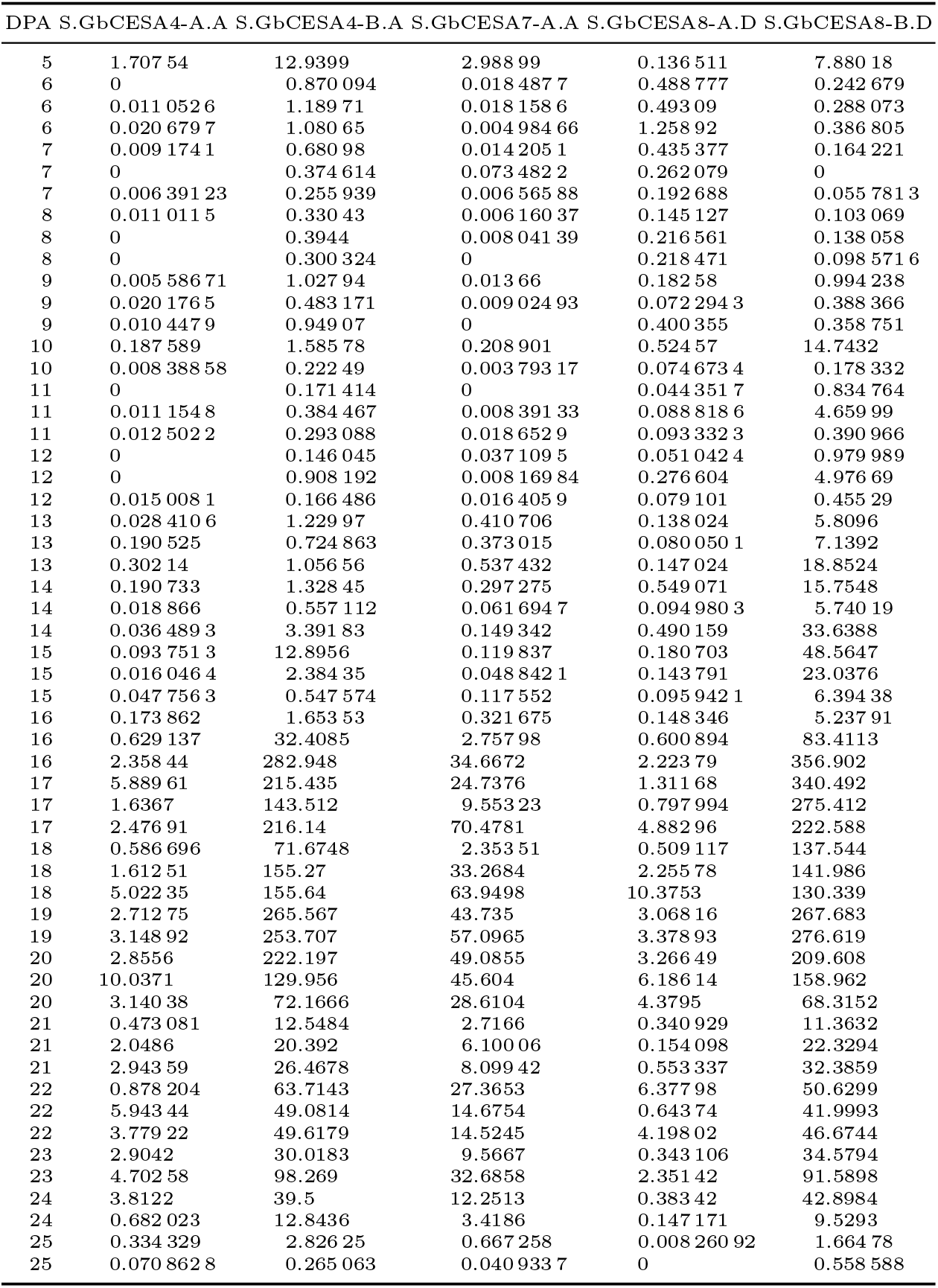
Transcriptomic data of Gossypium barbadense (Gb) in Transcripts Per Million (TPM) related to secondary cell wall cellulose synthesis. This dataset emphasizes the significant contributions of CESA4, CESA7, and CESA8 in cellulose biosynthesis.

**Table 3.**
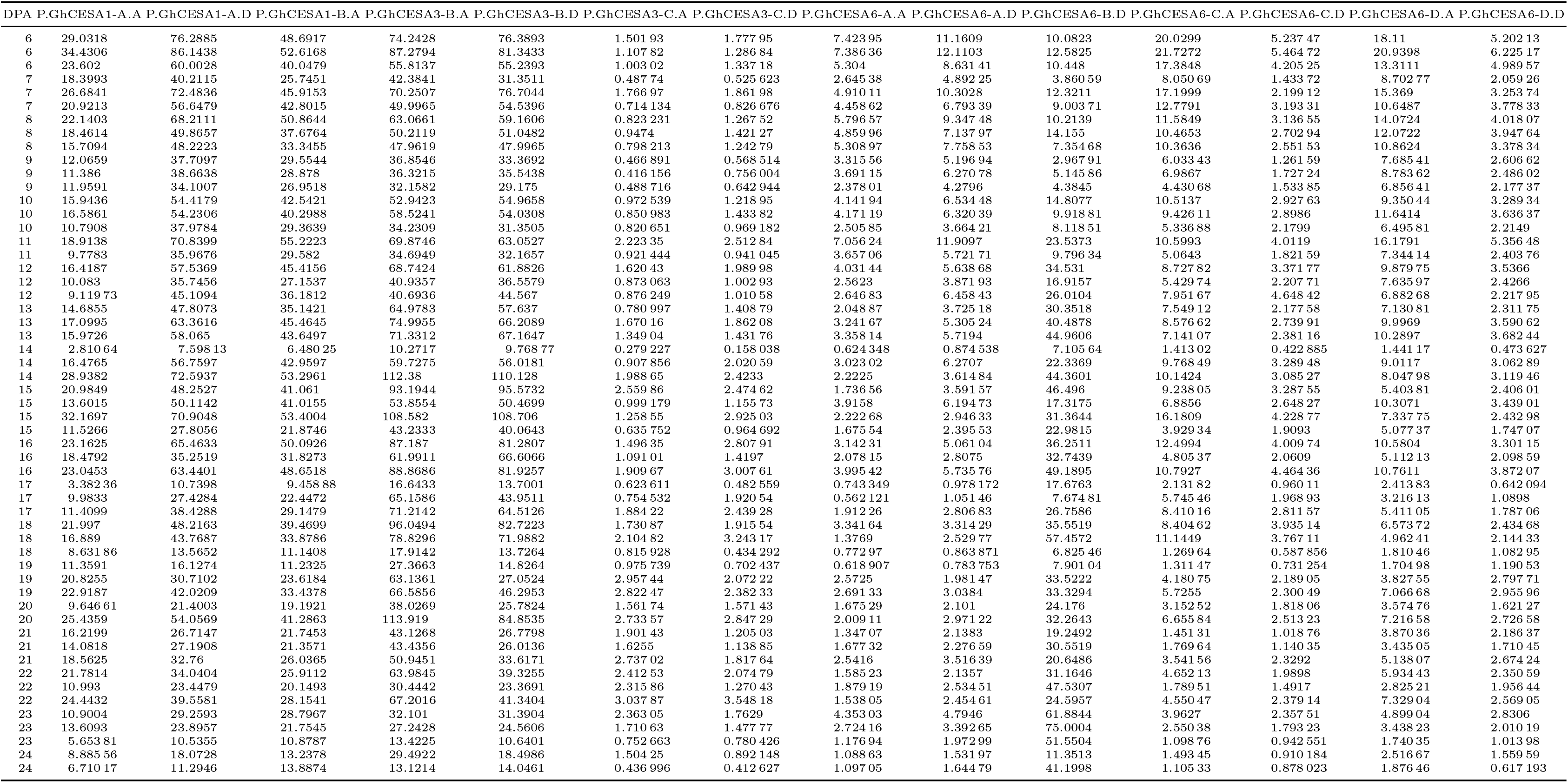
Transcriptomic data of *Gossypium hirsutum* (Gh) in Transcripts Per Million (TPM) related to primary cell wall cellulose synthesis. This dataset emphasizes the significant contributions of CESA1, CESA3, and CESA6 in cellulose biosynthesis.

**Table 4.**
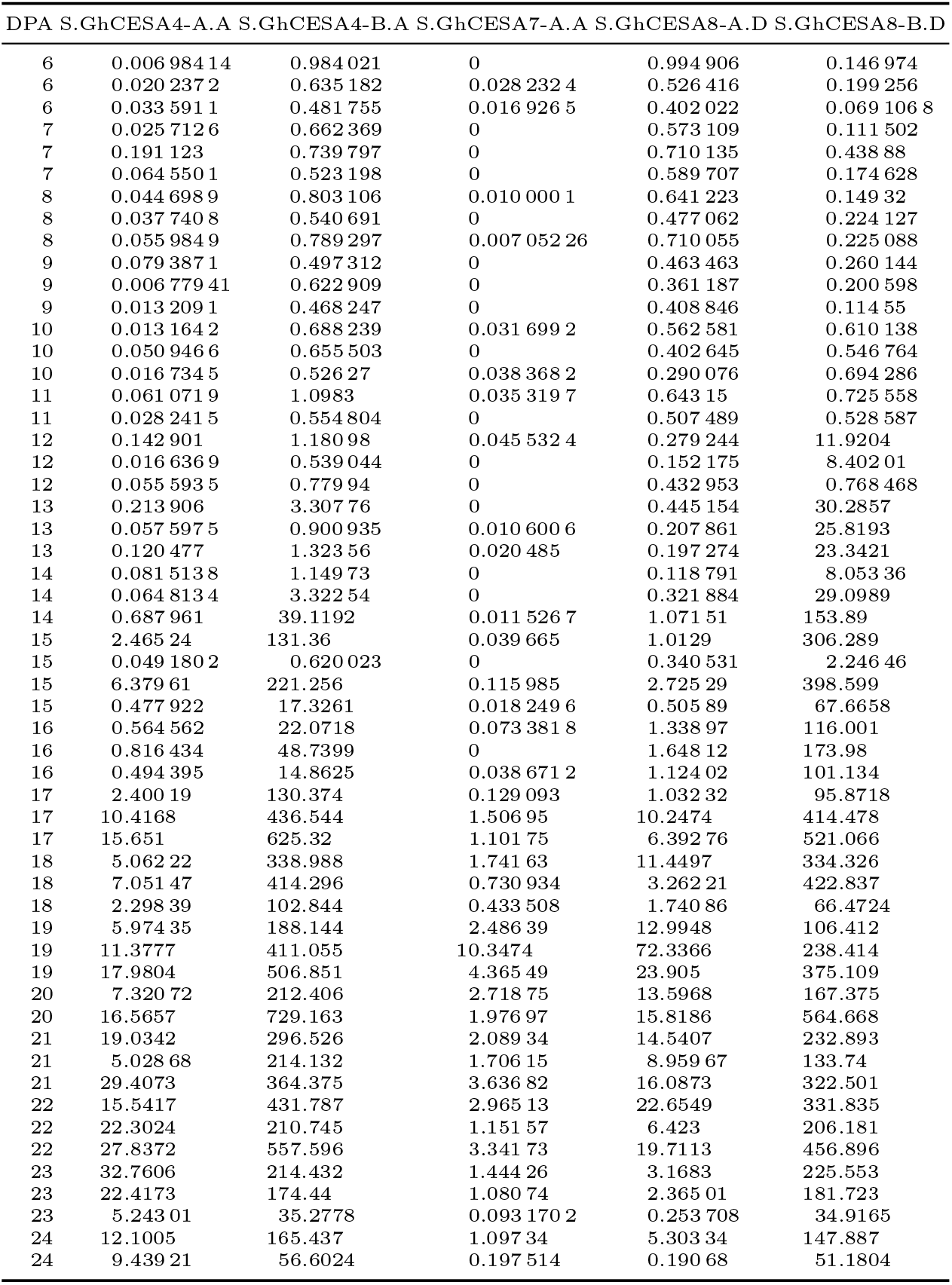
Transcriptomic data of *Gossypium hirsutum* (Gh) in Transcripts Per Million (TPM) related to secondary cell wall cellulose synthesis. This dataset highlights the expression levels of CESA4, CESA7, and CESA8 genes across different DPA values.

| DPA | S.GhCESA4-A-A | S.GhCESA4-B-A | S.GhCESA7-A-A | S.GhCESA8-A-D | S.GhCESA8-B-D |
| --- | --- | --- | --- | --- | --- |
| 6 | 0.00698414 | 0.984021 | 0 | 0.994906 | 0.146974 |
| 6 | 0.0202372 | 0.635182 | 0.0282324 | 0.526416 | 0.199256 |
| 6 | 0.0335911 | 0.481755 | 0.0169265 | 0.402022 | 0.0691068 |
| 7 | 0.0257126 | 0.662369 | 0 | 0.573109 | 0.111502 |
| 7 | 0.191123 | 0.739797 | 0 | 0.710135 | 0.43888 |
| 7 | 0.0645501 | 0.523198 | 0 | 0.589707 | 0.174628 |
| 8 | 0.0446989 | 0.803106 | 0.0100001 | 0.641223 | 0.14932 |
| 8 | 0.0377408 | 0.540691 | 0 | 0.477062 | 0.224127 |
| 8 | 0.0559849 | 0.789297 | 0.00705226 | 0.710055 | 0.225088 |
| 9 | 0.0793871 | 0.497312 | 0 | 0.463463 | 0.260144 |
| 9 | 0.00677941 | 0.622909 | 0 | 0.361187 | 0.200598 |
| 9 | 0.0132091 | 0.468247 | 0 | 0.408846 | 0.11455 |
| 10 | 0.0131642 | 0.688239 | 0.0316992 | 0.562581 | 0.610138 |
| 10 | 0.0509466 | 0.655503 | 0 | 0.402645 | 0.546764 |
| 10 | 0.0167345 | 0.52627 | 0.0383682 | 0.290076 | 0.694286 |
| 11 | 0.0610719 | 1.0983 | 0.0353197 | 0.64315 | 0.725558 |
| 11 | 0.0282415 | 0.554804 | 0 | 0.507489 | 0.528587 |
| 12 | 0.142901 | 1.18098 | 0.0455324 | 0.279244 | 11.9204 |
| 12 | 0.0166369 | 0.539044 | 0 | 0.152175 | 8.40201 |
| 12 | 0.0555935 | 0.77994 | 0 | 0.432953 | 0.768468 |
| 13 | 0.213906 | 3.30776 | 0 | 0.445154 | 30.2857 |
| 13 | 0.0575975 | 0.900935 | 0.0106006 | 0.207861 | 25.8193 |
| 13 | 0.120477 | 1.32356 | 0.020485 | 0.197274 | 23.3421 |
| 14 | 0.0815138 | 1.14973 | 0 | 0.118791 | 8.05336 |
| 14 | 0.0648134 | 3.32254 | 0 | 0.321884 | 29.0989 |
| 14 | 0.687961 | 39.1192 | 0.0115267 | 1.07151 | 153.89 |
| 15 | 2.46524 | 131.36 | 0.039665 | 1.0129 | 306.289 |
| 15 | 0.0491802 | 0.620023 | 0 | 0.340531 | 2.24646 |
| 15 | 6.37961 | 221.256 | 0.115985 | 2.72529 | 398.599 |
| 15 | 0.477922 | 17.3261 | 0.0182496 | 0.50589 | 67.6658 |
| 16 | 0.564562 | 22.0718 | 0.0733818 | 1.33897 | 116.001 |
| 16 | 0.816434 | 48.7399 | 0 | 1.64812 | 173.98 |
| 16 | 0.494395 | 14.8625 | 0.0386712 | 1.12402 | 101.134 |
| 17 | 2.40019 | 130.374 | 0.129093 | 1.03232 | 95.8718 |
| 17 | 10.4168 | 436.544 | 1.50695 | 10.2474 | 414.478 |
| 17 | 15.651 | 625.32 | 1.10175 | 6.39276 | 521.066 |
| 18 | 5.06222 | 338.988 | 1.74163 | 11.4497 | 334.326 |
| 18 | 7.05147 | 414.296 | 0.730934 | 3.26221 | 422.837 |
| 18 | 2.29839 | 102.844 | 0.433508 | 1.74086 | 66.4724 |
| 19 | 5.97435 | 188.144 | 2.48639 | 12.9948 | 106.412 |
| 19 | 11.3777 | 411.055 | 10.3474 | 72.3366 | 238.414 |
| 19 | 17.9804 | 506.851 | 4.36549 | 23.905 | 375.109 |
| 20 | 7.32072 | 212.406 | 2.71875 | 13.5968 | 167.375 |
| 20 | 16.5657 | 729.163 | 1.97697 | 15.8186 | 564.668 |
| 21 | 19.0342 | 296.526 | 2.08934 | 14.5407 | 232.893 |
| 21 | 5.02868 | 214.132 | 1.70615 | 8.95967 | 133.74 |
| 21 | 29.4073 | 364.375 | 3.63682 | 16.0873 | 322.501 |
| 22 | 15.5417 | 431.787 | 2.96513 | 22.6549 | 331.835 |
| 22 | 22.3024 | 210.745 | 1.15157 | 6.423 | 206.181 |
| 22 | 27.8372 | 557.596 | 3.34173 | 19.7113 | 456.896 |
| 23 | 32.7606 | 214.432 | 1.44426 | 3.1683 | 225.553 |
| 23 | 22.4173 | 174.44 | 1.08074 | 2.36501 | 181.723 |
| 23 | 5.24301 | 35.2778 | 0.0931702 | 0.253708 | 34.9165 |
| 24 | 12.1005 | 165.437 | 1.09734 | 5.30334 | 147.887 |
| 24 | 9.43921 | 56.6024 | 0.197514 | 0.19068 | 51.1804 |

**Table 5.** Performance Comparison of Regression Models Using AFM Features (Mean (Std))

| Target | Model | $R^2$ | MAE | MSE |
| --- | --- | --- | --- | --- |
| Boll Length (cm) | LinearRegression | 0.76 (0.05) | 0.37 (0.02) | 0.23 (0.03) |
|  | Ridge | 0.76 (0.04) | 0.37 (0.02) | 0.23 (0.03) |
|  | Lasso | 0.76 (0.04) | 0.37 (0.02) | 0.23 (0.03) |
|  | ElasticNet | 0.76 (0.04) | 0.38 (0.02) | 0.23 (0.03) |
|  | RandomForest | 0.79 (0.08) | 0.31 (0.05) | 0.20 (0.07) |
| Cellulose Content<br>(mg/cotton boll) | LinearRegression | 0.71 (0.05) | 0.34 (0.03) | 0.29 (0.07) |
|  | Ridge | 0.71 (0.04) | 0.34 (0.03) | 0.29 (0.07) |
|  | Lasso | 0.71 (0.04) | 0.34 (0.03) | 0.29 (0.06) |
|  | ElasticNet | 0.72 (0.04) | 0.34 (0.04) | 0.28 (0.07) |
|  | RandomForest | 0.92 (0.02) | 0.16 (0.03) | 0.07 (0.03) |

| Target | Model | $R^2$ | MAE | MSE |
| --- | --- | --- | --- | --- |
| Boll Length (cm) | LinearRegression | 0.85 (0.04) | 0.27 (0.02) | 0.14 (0.03) |
|  | Ridge | 0.85 (0.07) | 0.27 (0.04) | 0.14 (0.05) |
|  | Lasso | 0.85 (0.07) | 0.27 (0.05) | 0.13 (0.05) |
|  | ElasticNet | 0.86 (0.06) | 0.26 (0.04) | 0.13 (0.05) |
|  | RandomForest | 0.87 (0.09) | 0.24 (0.05) | 0.11 (0.07) |
| Cellulose Content<br>(mg/cotton boll) | LinearRegression | 0.74 (0.07) | 0.33 (0.06) | 0.26 (0.08) |
|  | Ridge | 0.75 (0.06) | 0.32 (0.04) | 0.25 (0.07) |
|  | Lasso | 0.75 (0.06) | 0.32 (0.05) | 0.25 (0.07) |
|  | ElasticNet | 0.75 (0.05) | 0.31 (0.04) | 0.25 (0.07) |
|  | RandomForest | 0.86 (0.09) | 0.18 (0.06) | 0.14 (0.10) |

### 1.2. Mechanics Models in PeakForce QNM Analysis

The force-displacement data were fitted using the Hertz model (spherical indenter) or the Sneddon model (conical indenter) in order to calculate Young’s modulus. The equation of the force-displacement relation for the Hertz model is [1]:

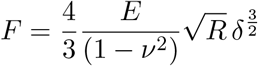

where *F* is the applied force, *E* is Young’s modulus, *ν* is Poisson’s ratio, *δ* represents the indentation depth, and *R* is the radius of the spherical indenter.

For conical indenters, the Sneddon model is used, where the force is expressed as:

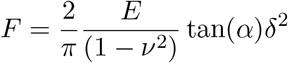

Here, *E* represents Young’s modulus, *ν* is Poisson’s ratio, *α* is the half-angle of the conical indenter, and *δ* is the indentation depth.

The Hertz model is applicable to the case when the AFM probe is terminated by a spherical tip, and the Sneddon model to the case when the probe is conical. Both models operate with an infinitely stiff AFM tip interacting with an elastic sample or cylindrical nanostructure. Although both models accurately describe the indentation mechanics, absolute values of the Young’s modulus may differ based on the chosen model, although general trends are identical. Figure S4 shows the process used to calculate Young’s modulus in this study. NanoScope Analysis software was employed to analyze .pfc files obtained from PeakForce QNM AFM experiments (producing high-resolution AFM height and peak force error images as well as high-resolution mechanical property maps). Multiple points along the microfibrils were selected to extract corresponding force–distance curves. These curves were then processed using the Indentation feature of NanoScope Analysis software, where the Hertz model was applied to determine Young’s modulus. This approach enables localized and accurate nanomechanical characterization of fiber surfaces.

### 1.3. MATLAB Processing and Analysis

In this section, we provided a detailed explanation of the methods used to quantify angles crossover, crossover counts, and intensity measurements in microfibrils structures using MATLAB. These analytical approaches are essential for assessing fiber alignment and structural characteristics in our samples. Additionally, we included example MATLAB code and a step-by-step guide for implementing these methods on similar datasets.

#### Angles of Crossover

To determine the angles crossover, we developed a MATLAB script that overlays two adjustable lines onto fiber strands to align with observed crossovers. This method allowed for precise measurement of fiber orientation at intersection points. As illustrated in Figure S5(a), we analyzed a selected region from a 8 DPA Gh fiber sample. The MATLAB script enabled users to manipulate three control points along the two lines, allowing for accurate positioning over the desired crossover locations. The script relies on the Image Processing Toolbox in MATLAB, which is required for image handling and visualization. The following MATLAB code generates two adjustable lines over an image, allowing users to align them with fiber crossovers. To use this functionality, images must be stored in a location accessible to MATLAB and saved in a file format supported by the Image Processing Toolbox. When the code is executed, the selected image appears on the screen with two interactive lines. Users can adjust the position of three control points along the lines, enabling precise alignment with fiber structures and facilitating angle measurements.

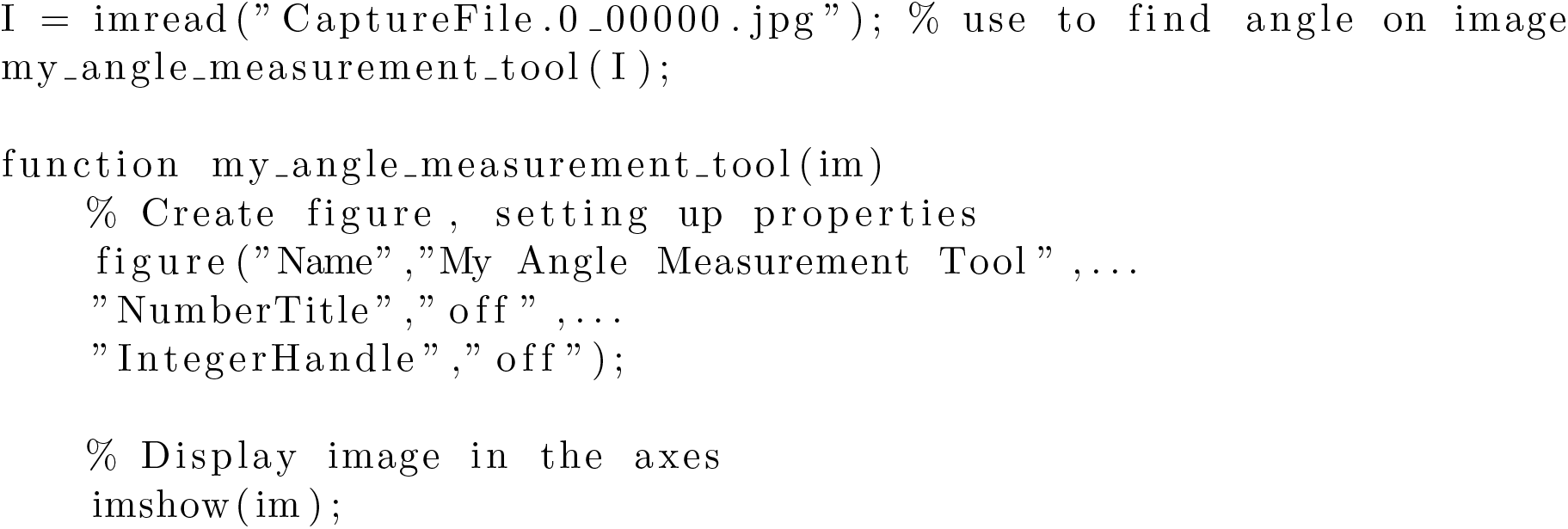
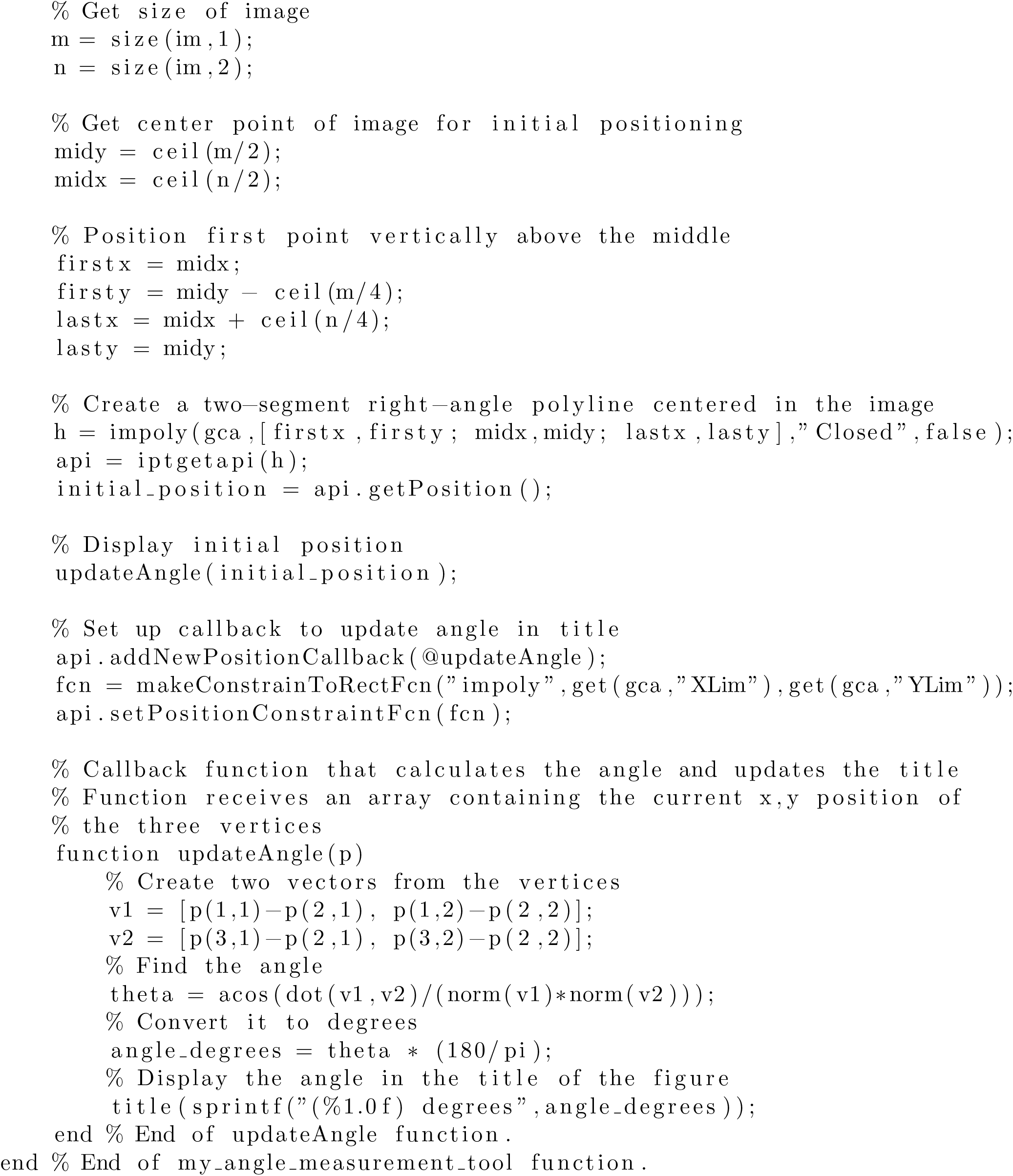

#### Crossover count

For our data on crossover count, we used NanoScope Analysis 3.00 to obtain crossover count data across different days for Gh and Gb. To collect this data, we opened .spm files into NanoScope Analysis, created a 500 nm × 500 nm box over a selected region in the image, and counted the number of microfibril crossovers.

Using the Peak Force Error view in NanoScope Analysis, we utilized the ‘Crop’ and ‘Split’ tools to define a selected area within an image. By using the Snipping Tool in Windows, we extracted that section and manually marked each crossover within the selected region that is showed in Figure S5(b). The number of crossovers is recorded for four images per day for both Gh and Gb.

#### Intensity

To obtain intensity measurements of our samples, we used MATLAB code to convert sample images into grayscale and then measured intensity along a user-defined line. Figure S5(c) and S5(d) below provide examples of how the code was used to extract intensity measurements.

The following code was used to obtain intensity measurements. To generate a graph of the results, the image file must be compatible with the Image Processing Toolbox in MATLAB. When the code is executed, the selected image appears on the screen. Users can left-click and drag to draw a line along a chosen section of the image. Right-clicking generates a graph displaying the corresponding intensity measurements. Additionally, multiple line segments can be created by drawing the first segment and left-clicking again to add another.

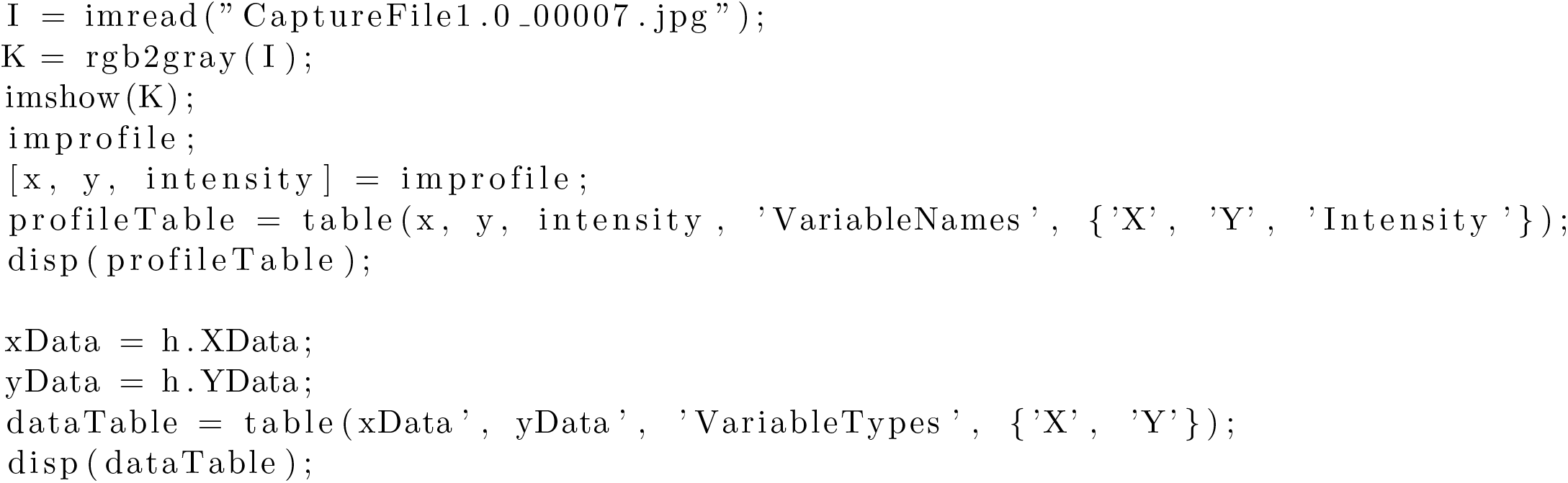

### 1.4 Polysaccharides impacting the thickness of microfibrils

Monosaccharide analyses of cotton fiber cell walls demonstrated the presence of a large amount of glucose, mostly coming from such amorphous glucan, which gradually increased between 6-22 DPA and then maintained at the same level up to 25 DPA [2]. At the same time, individual cellulose microfibrils bundle together into larger fibrillar structures, which are typical in cotton and other plant fibers. It was demonstrated by Cosgrove et al. [3] that during synthesis, individual cellulose chains usually coalesce into a protofibrils of three chains, followed by coalescence of the latter into cellulosic microfiber of 18-24 chains. Later, such individual microfibrils bundle into higher-order larger microfibrils in the native walls. These can be observed using high-resolution nanoscale imaging technologies and these factors can promote the self-assembling of individual cellulosic microfibrils into higher-order bundles, particularly promoting the transition to the secondary cell wall cellulosic microfibril formation [4, 3]. This occurs rapidly in cotton fibers and reduction in matrix polysaccharide content during fiber development can dynamically modulate the interaction of adjacent cellulosic microfibrils. That is why at 8 and 12 DPA, the obtained images show microfibrils with lower dimensions, likely due to larger coverage by hemicellulosic polysaccharides. On the other hand, at 22 DPA, when cellulose biosynthesis dramatically increases and matrix polysaccharide relative content drops significantly, the bundling of cellulose microfibrils into higher-order microfibrils with more parallel orientation takes place, likely due to the increased number and density of the secondary cell walls cellulose-synthesizing complexes (CSCs) on cotton fiber membrane.

### 1.5 Feature Selection For Predictive Model

In this study, we employed a two-step feature selection approach to ensure that only informative variables were included in our machine learning models.

#### Data Distribution Assessment via Gaussian Process Regression (GPR)

We utilized Gaussian Process Regression (GPR) to visualize the data distributions of individual transcriptomic features. GPR is a non-parametric method that defines a probability distribution over potential functions that fit the observed data, assuming that any collection of points follows a multivariate Gaussian distribution. Features that did not exhibit distinct, informative patterns were considered non-informative and were excluded from further analysis. As shown in Figure S6, several transcriptomic features displayed distributions lacking significant structure, suggesting that their inclusion could introduce noise and adversely affect model accuracy and generalization.

#### Redundancy Reduction through Correlation Analysis

To further refine our feature set, we computed pairwise Pearson correlation coefficients among the transcriptomic features. The initial heatmap in Figure S7 reveals clusters of highly correlated features, indicating potential multicollinearity. We applied a correlation threshold (*r >* 0.97) to remove redundant features, retaining one representative gene from each highly correlated cluster as identified through hierarchical clustering. The resulting optimized heatmap (Figure S8) shows a significant reduction in redundant correlations, thereby preserving biological signal while minimizing noise.

### 1.6. Predictive Models and Outcome

In the main text, we demonstrated that combining transcriptomic (CESAs) and nanomechanical (AFM) features yields superior predictive performance for both macroscale phenotypic traits: boll length and cellulose content. To provide a comprehensive view of each individual feature set’s contribution, Tables 5 and 6 summarize the predictive outcomes when models were trained solely on (1) nanomechanical features (AFM-only) or (2) transcriptomic features (CESAs-only), respectively.

Specifically, Table 6 highlights the performance of regression models trained with only AFM-derived nanomechanical measurements (e.g., width, height, roughness, Young’s modulus, angle, and crossover number). Table 5, in turn, focuses on models trained solely with transcriptomic data (primary and secondary wall CESA-encoding genes). Each table reports standard regression metrics—*R*^2^, MAE, and MSE—across multiple models (LinearRegression, Ridge, Lasso, ElasticNet, and RandomForest), using 5-fold cross-validation. All metrics are presented as mean (standard deviation) and rounded to three decimal places for clarity.

While these single-modality approaches show moderate predictive power, the results underscore the limitations of using either CESA-only or AFM-only features. As discussed in the main text (Results and Discussion), combining transcriptomic and nanomechanical features (CESAs+AFM) significantly enhances model accuracy. This improvement further highlights the complementary nature of gene expression profiles and nano-scale morphological traits in capturing the complexity of cotton fiber development. Our proposed approach showcased the importance of ‘nano-omics’ for the first time for macroscale plant phenotypic feature prediction. This will add a new dimension to the molecular breeding process, making it faster and more informed, eventually benefiting plant science researchers and farmers.

